# Learning shapes neural codes for sensory-motor integration in the tail of the striatum

**DOI:** 10.64898/2026.05.05.722944

**Authors:** Ivan Linares-Garcia, Sofia E. Juliani, Jessie Yi, Diego Castro, David J. Margolis

## Abstract

Separating meaningful sensory stimuli from irrelevant ones requires learning sensorimotor associations, but how sensory-linked striatal circuits acquire and maintain these associations is unclear. We longitudinally imaged direct- and indirect-pathway (D1 and A2a) spiny projection neurons (SPNs) in the tail of the striatum (TS) as mice learned to push or pull a joystick in response to auditory cues in either a stimulus-response association (go/omit) task or a two-alternative forced choice (2AFC) task. Learning in both tasks increased the fraction and strength of task-modulated TS SPNs across the sound, action, and reward epochs, yet individual neuron selectivity often switched over days between behavioral epochs. In spite of individual neuron variability, population activity of direct and indirect pathways became aligned with characteristic behavioral features during learning: D1-SPNs dominated the ‘action’ category, A2a-SPNs were biased toward the ‘mixed’ category (multiple epochs), and both SPN types showed ‘sound’ category specificity that depended on the sound-action association. Trial-wise modeling revealed a reweighting of behavioral predictors within the action window, with reward gaining and movement losing predictive weight. Learning the two-choice task led to a higher prevalence of association-preferring neurons and better behavioral decoding within the sound window than in the action/reward window, reflecting a task-dependent prioritization of sensory information. Association-preferring neurons also showed a stable local distance-similarity relationship, with nearby neurons more similar than distant neurons across learning. Together, our results support a population mechanism in TS during learning in which neurons from both direct and indirect pathways are recruited and take on distinct behavioral roles that vary with performance and task complexity.

## Introduction

Throughout life, we are continuously faced with sensory stimuli, and we must learn through experience to assign meaning to them in order to guide appropriate actions (Makino et al., 2016). Sensorimotor associative learning is therefore fundamental for survival, for example, understanding the meaning of a red or green traffic light, or, for a mouse, distinguishing the sound of a mate’s approach from the slither of a predator. Yet how the brain implements sensorimotor associations remains incompletely understood.

The striatum, as the major input nucleus of the basal ganglia, is central to learning and the adaptive control of behavior. Dorsal striatal circuits regulate motor sequences, goal-directed actions, and stimulus-response associations (Redgrave et al., 1999; Kreitzer and Malenka, 2008; Gerfen and Surmeier, 2011; Valjent and Gangarossa, 2021). Striatal activity and plasticity have been linked to motor learning and skill refinement (Hwang et al., 2019, 2021), and reinforcement learning (Panigrahi et al., 2015; Yttri and Dudman, 2016; Roth et al., 2024). Spiny projection neurons (SPNs; also known as medium spiny neurons, MSNs) mediate the direct-and indirect-pathway outputs of the striatum, corresponding broadly to D1-SPNs and D2/A2a-SPNs, respectively. Classic antagonistic models emphasize direct-pathway facilitation and indirect-pathway suppression of actions (Albin et al., 1989; DeLong, 1990; Kravitz et al., 2010), whereas concurrent-activation models emphasize co-recruitment of both pathways around movement onset (Cui et al., 2013; Tecuapetla et al., 2014). A selection and inhibition framework helps reconcile these observations by proposing that both pathways act together to enable selected actions while suppressing competing motor programs (Mink, 1996; Yttri & Dudman, 2016; Oldenburg & Sabatini, 2015; Sheng et al., 2019; Cruz et al., 2022). SPN population activity can also show functional spatiotemporal organization during learned behavior, and striatal activity can reflect topographically organized cortical activity (Barbera et al., 2016; Klaus et al., 2017; Parker et al., 2018; Peters et al., 2021).

Much of what we know about striatal learning comes from motor control and action sequencing, but the striatum also participates in sensory learning when sensory evidence must be translated into action. Neurons in dorsolateral striatum (DLS) receive robust sensory inputs across modalities (Hunnicutt et al., 2016). Glutamatergic drive from sensory cortex and thalamus shapes SPN activity in behaviorally relevant ways, and dopamine further modulates these inputs during learning (Lee et al., 2019; Ponvert & Jaramillo, 2019; Hidalgo-Balbuena et al., 2019; Sanabria et al., 2024; Schultz et al., 1997; Hollerman & Schultz, 1998). In whisker-based tasks, direct-pathway SPNs show early sensory responses, and their activation can substitute for sensory stimulation to trigger the behavioral response (Sippy et al., 2015).

The posterior tail of the striatum (TS) has emerged as a sensory-linked striatal sector with strong sensory input, including prominent auditory input (Chen et al., 2019), visual input (Hunnicutt et al., 2016), weaker coupling to overall movement compared to the anterior portion(Guo et al., 2018), and specialized dopamine signaling biased toward stimulus novelty and intensity (Menegas et al., 2018; Green et al., 2024). Recent work further shows that movement-related dopamine activity in TS can encode an action prediction error that serves as a value-free teaching signal to reinforce stable sound-action associations (Greenstreet et al., 2025). In TS, stable sound codes can support decisions (Guo et al., 2018), parvalbumin (PV) interneurons regulate sensory responses and auditory-guided behavior (Li et al., 2024), pathway-specific choice signals have been observed during auditory tasks (Tang et al., 2025), and indirect-pathway neurons can regulate inhibitory control over sensory-driven behavior (Ferrigno et al., 2025). Auditory representations in posterior striatum can also emerge rapidly during associative learning (Znamenskiy & Zador, 2013; Guo et al., 2018; Xiong et al., 2015). Together, these findings suggest that sensory-linked striatal circuits can rapidly acquire task-relevant stimulus codes as animals learn cued actions in response to sound. However, several key questions remain unresolved. We still lack a clear picture of how TS population activity is reorganized through learning as sounds acquire behavioral meaning and become linked to specific actions, and how this reorganization differs across direct and indirect pathway SPNs. It is also unclear whether learned TS representations are maintained in a stable form over days or instead reconfigured as learning progresses, including whether any functional spatiotemporal organization emerges within TS. Finally, beyond stimulus-response behavior, it remains unclear how TS activity supports within-session selection between opposing actions based on sound identity, and how such representations change with learning.

Here we addressed these gaps by longitudinally imaging pathway-identified TS SPNs as mice learned sound-action associations. We first used a stimulus-response (go/omit) task to ask how learning recruits TS neurons across sound, action, and reward epochs, whether this recruitment remains stable across days, and how direct and indirect pathway populations contribute as these associations are learned. We then used a two-choice (2AFC) task, in which two sounds were interleaved within the same session and mice selected between push and pull choices on every trial based on sound identity, to ask how TS activity differentiates the two tone-action associations within a session and how these signatures change with learning. Learning increased both the fraction and strength of task modulation in TS SPNs.

Across days, the overall distribution of neurons across functional categories remained stable even as the identities of contributing neurons shifted, consistent with stable population-level output and drifting single-cell participation. Recruitment was learning and association-sensitive: action-related signals became increasingly D1-dominated, Mixed neurons (neurons active across multiple task epochs), were biased toward A2a, and sound-window pathway biases depended on the learned association. Trial-wise GLM modeling revealed a learning-related reweighting during the action epoch, with reward-linked predictors gaining influence as movement-linked predictors diminished. In the two-choice task, association-preferring activity and decoding were strongest in the sound window, increased with learning, and dropped after reversal. Together, these findings support a view of TS as a learning-related striatal substrate in which both pathways are concurrently recruited but assume distinct, reweighted roles as sound-action behavior is learned.

## Results

We imaged TS SPNs in head-fixed mice as they learned tone-guided joystick-based forelimb actions in an open-source auditory-motor task (Linares-Garcia et al., 2025; Fig. 1A,D). We performed longitudinal two-photon calcium imaging through a GRIN lens while mice expressed GCaMP8f, allowing us to track large TS populations across learning. Using double-transgenic D1-Cre × Ai14 and A2a-Cre × Ai14 mice, we identified pathway populations by tdTomato co-expression, assigning tdTomato+GCaMP (“yellow”) neurons to the Cre-defined pathway and treating GCaMP-only neurons as putative neurons of the opposite pathway (Fig. 1B).

**Figure 1.**
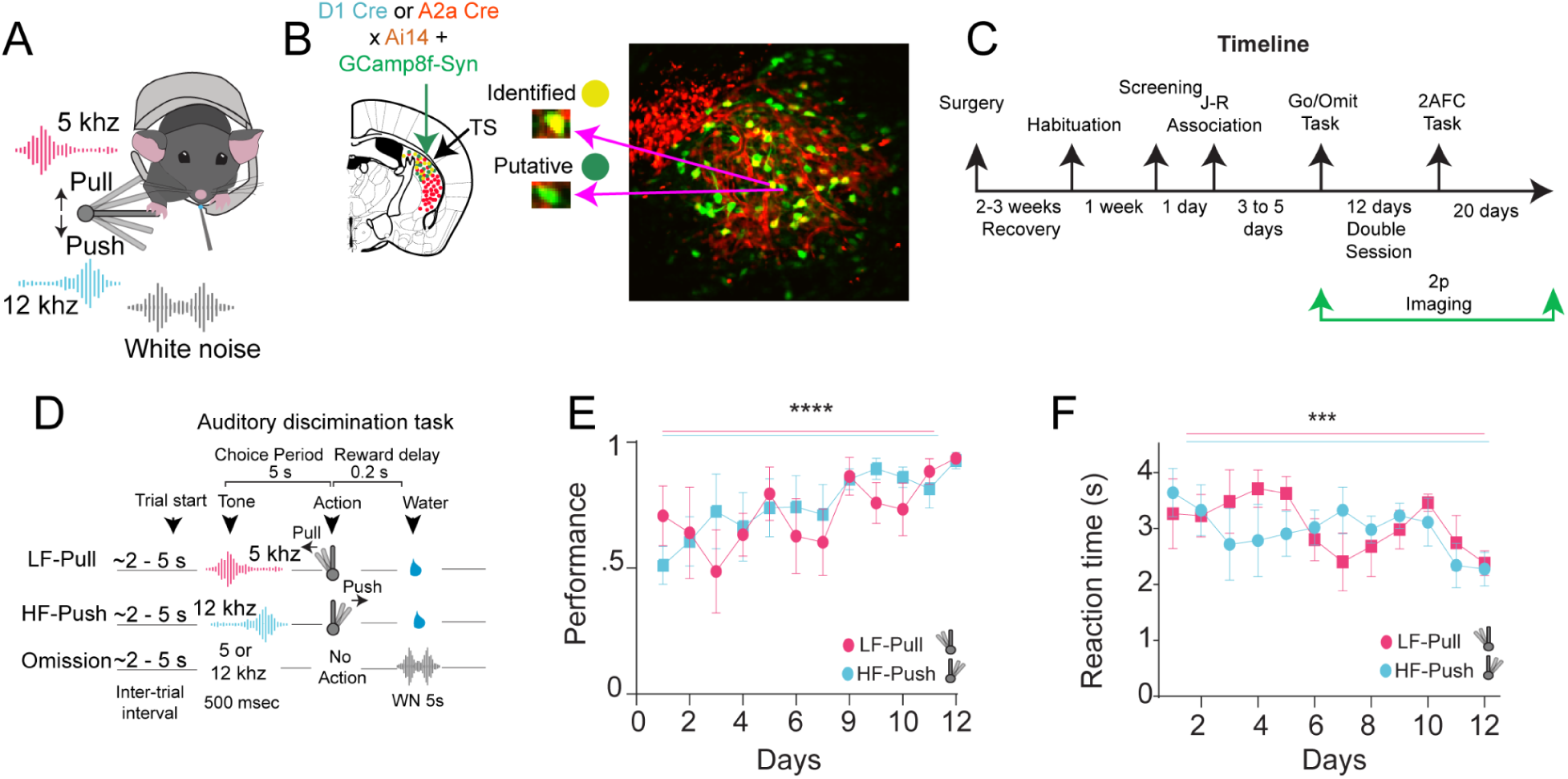
Behavioral training on the Go/Omit task during longitudinal TS imaging. **A,** Schematic of the head-fixed joystick task. Mice used the right paw to displace a joystick in response to sounds to obtain water reward. **B,** Example two-photon field of view in TS through a GRIN lens. Neurons expressing GCaMP8f are shown in green; tdTomato+GCaMP8f neurons (“yellow”) identify Cre-defined pathway populations in D1-Cre × Ai14 or A2a-Cre × Ai14 mice. **C,** Experimental timeline from surgery and recovery through habituation, shaping, joystick–reward association (J–R), longitudinal imaging during the stimulus–response (go/omit) task, and the two-choice (2AFC) task. **D,** Trial structure for the single-association auditory discrimination task. After a random inter-trial interval (2-5 s), a 500-ms tone was presented, followed by a 500-ms grace period and a 5-s response window. Correct joystick displacements yielded water reward with 90% probability (10% white noise, WN). Omissions triggered 5 s of white noise before the next trial. Training was performed in separate sessions for the low-frequency pull (LF-Pull) and high-frequency push (HF-Push) associations. **E,** Performance (correct / (correct + omissions)) increased across 12 days of training for both associations (Kruskal-Wallis test across days, p < 0.05). **F,** Reaction time, time to reach the displacement threshold, decreased (faster responses) with learning from early to late training for both associations (Kruskal-Wallis test across days, p < 0.001).

We then used two complementary task designs. In a stimulus-response (go/omit) task, each session involved a single tone-action association and trials were scored as correct versus omission (Fig. 1C,D). In a two-choice (2AFC) task, both tones were interleaved within the same session and mice selected between push and pull on every trial based on sound identity, enabling within-session comparisons of neural activity for competing actions under the same behavioral context (Fig. 6A; Supplementary Fig. 5A).

In the stimulus-response (go/omit) task, mice learned two tone-guided associations trained in separate sessions: a high-frequency (HF; 12 kHz) tone cued a push, and a low-frequency (LF; 5 kHz) tone cued a pull (Fig. 1D). For both associations, performance (correct / (correct + omissions)) increased across 12 training days (Fig. 1E; Kruskal-Wallis, p < 0.05), and reaction times decreased with learning (Fig. 1F; Kruskal-Wallis, p < 0.001).

Across recorded sessions, we selected five representative days spanning low (∼50% incorrect trials) to high performance (∼90% correct trials) to examine how TS SPNs were modulated across task epochs (Supplementary Fig. 1D,E). For HF–Push, mean performance values (mean ± s.e.m.) across Sessions 1–5 were 0.58 ± 0.08, 0.71 ± 0.04, 0.84 ± 0.02, 0.91 ± 0.01, and 0.96 ± 0.02, respectively (Fig. 2B-D); for LF–Pull, they were 0.45 ± 0.05, 0.66 ± 0.05, 0.80 ± 0.03, 0.89 ± 0.02, and 0.96 ± 0.01 (Fig. 2F-H).

**Figure 2.**
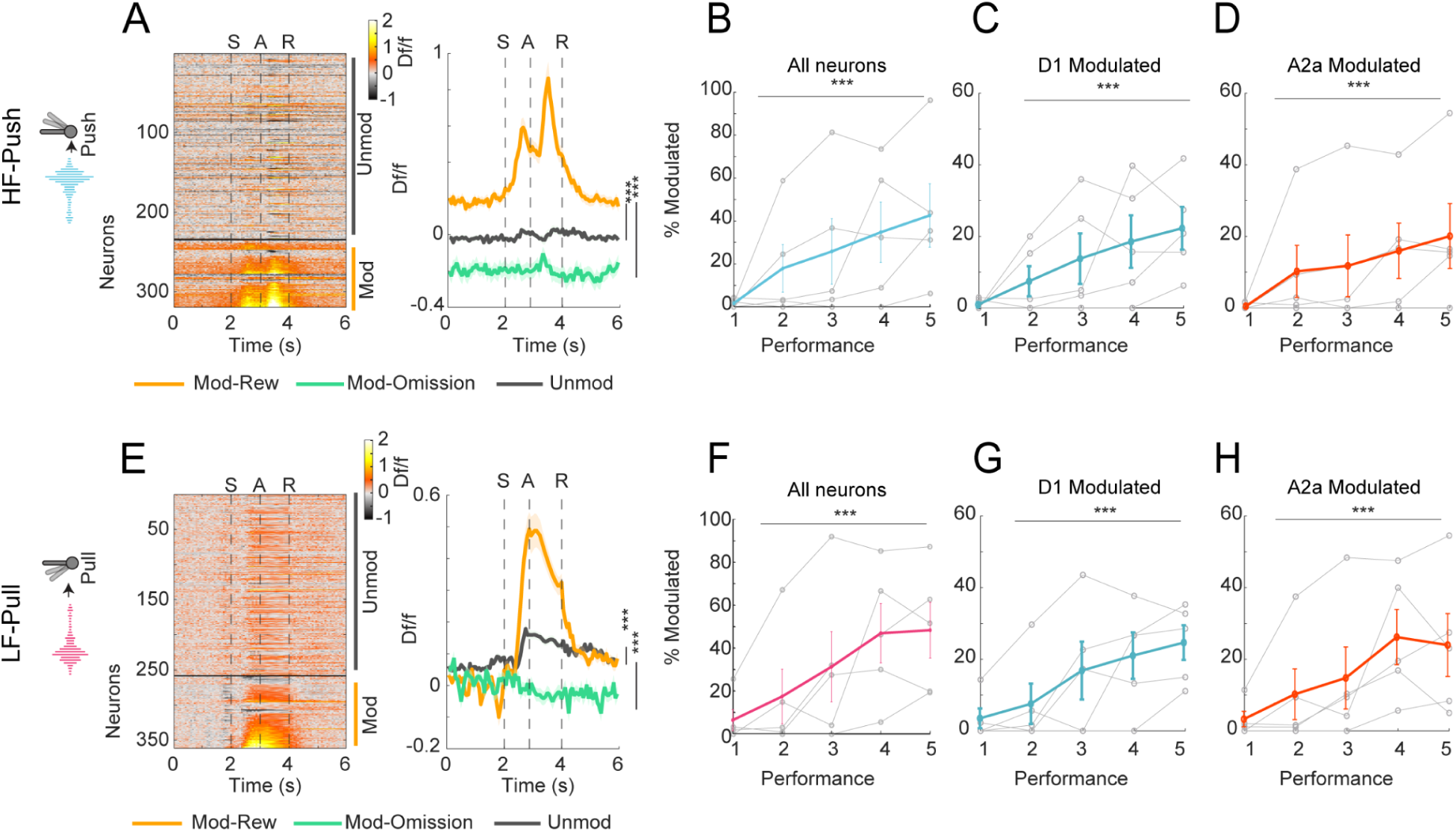
Task-modulated TS SPNs increase with learning across pathways. **A,** Example population activity on a high-performance day for the HF-Push association. Heatmap shows trial-averaged activity for all recorded TS SPNs, separated into task-modulated neurons (bottom) and unmodulated neurons (top). Right, mean activity traces for task modulated neurons on correctly rewarded trials (Mod-Rew), the same modulated neurons on omission trials (Mod-Omission), and unmodulated neurons on correct trials (Wilcoxon signed-rank test, p < 0.001). **B,** Fraction of task-modulated neurons across five representative sessions ranked from lowest to highest performance (Session 1 = lowest performance; Session 5 = highest performance; gray, individual animals; solid color, population summary; Kruskal–Wallis test, p < 0.05). **C-D,** Same as **B**, shown separately for D1 (**C**) and A2a (**D**) pathway populations (Kruskal-Wallis test across days, p < 0.05). **E-H,** Same as **A-D**, for the LF-Pull association.

A neuron was classified as modulated if its Area Under the Receiver Operating Characteristics (auROC) curve in any behavioral window (Sound/Action/Reward) exceeded 0.6 or fell below 0.4, with significance assessed using 1,000 label permutations across all trials for that neuron. On a high-performance day, the HF-Push association modulated TS SPNs showed stronger activity on rewarded trials compared with omitted trials from the same neurons, and compared with unmodulated neurons on rewarded trials, for both sound-action associations (Fig. 2A Wilcoxon signed-rank test, p < 0.001). The number of modulated neurons increased significantly with performance for the full population (Fig. 2B; Kruskal-Wallis across days, p < 0.05) and for both D1 and A2a subpopulations analyzed separately (Fig. 2C,D; Kruskal-Wallis across days, p < 0.05), with the same pattern in the LF-Pull association (Fig. 2E-H).

Together, these results show that learning progressively recruits TS SPNs across sound, action, and reward epochs, with correct trials eliciting stronger activity than omissions. Importantly, mean TS population activity showed no reliable zero-lag coupling to joystick displacement (Supplementary Fig. 1B,C), arguing against a unitary motor command and instead supporting a learning-dependent task representation in TS.

To determine how task-modulated TS SPNs were distributed across task epochs and how their tuning evolved with learning, we assigned neurons to four functional categories based on the epoch of their modulation (auROC with permutation significance). Neurons were labeled Sound, Action, or Reward if they were selectively modulated only in the sound (0-600 ms after tone onset), action (1 s, time-normalized to movement onset by linear resampling), or reward window (0-1 s after water release), respectively; neurons modulated in more than one window were labeled Mixed (Fig. 3A; HF-Push and Fig. 3E; LF-Pull).

**Figure 3.**
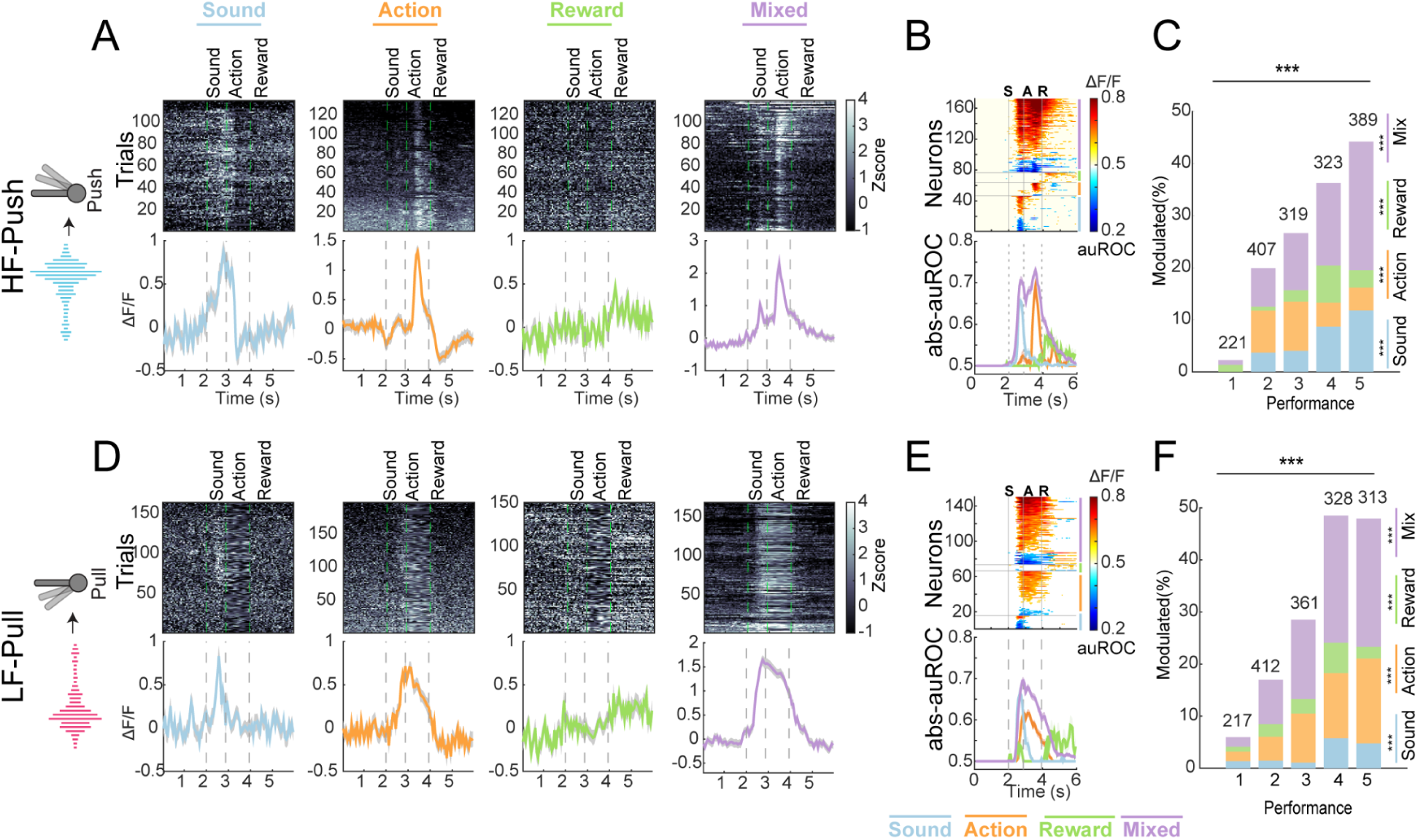
Behavioral categories expand with learning. **A,** HF-Push association. Example neurons from each auROC-defined category on a high-performance day, aligned to trial events: Sound (tone + grace period), Action (movement-aligned window), Reward (reward-aligned window), and Mixed (modulated in more than one window). **B,** HF-Push population responses by category. Top, heatmaps of absolute auROC values across neurons within each category. Bottom, mean activity traces for each category. **C,** HF-Push category composition across learning/performance days. Stacked fractions of Sound, Action, Reward, Mixed, and Unmodulated neurons across five representative sessions (χ² test of independence across days; see Results). **D-F,** Same as A-C for the LF-Pull association. **D,** example category neurons; **E,** population responses by category; **F,** category composition across days (χ² test; see Results).

Neurons populated all four categories for both sound-action associations (Fig. 3B,F). Within a given day, the distribution across categories changed with performance (Fisher’s exact test, p < 0.05), with higher-performance days showing more Sound, Action, and Mixed tuned neurons (Fig. 3C,I). In both sound-action associations, the distribution of categories changed significantly with performance (HF-Push: χ²(16) = 218.5, p < 0.001; LF-Pull: χ²(16) = 210.4, p < 0.001; Fig. 3C,G). In HF-Push, the fractions of Sound, Reward, and especially Mixed neurons increased across days (all p ≤ 0.0001), while Action neurons showed a significant but non-monotonic profile (p = 0.0001). In LF-Pull, Action and Mixed neurons expanded most strongly with performance (both p ≤ 0.0001), with more modest but significant changes in Sound and Reward (p ≤ 0.01). Thus, higher-performance days contain a larger proportion of Sound-, Action-, and Mixed-tuned neurons, with the largest dynamic range in the Mixed category (Fig. 3F).

We then tested whether category tuning changed with learning. For each neuron, we defined peak tuning as the maximum absolute auROC value within its category window (higher |auROC| indicates stronger tuning relative to baseline). Overall, peak tuning increased with performance in the HF-Push association for Sound, Action, and Mixed neurons (Kruskal-Wallis test across days, p < 0.05; Fig. 4A), while peak tuning increased with performance in the LF-Pull association for Action and Mixed neurons (Kruskal-Wallis test across days, p < 0.05; Fig. 4D).

**Figure 4.**
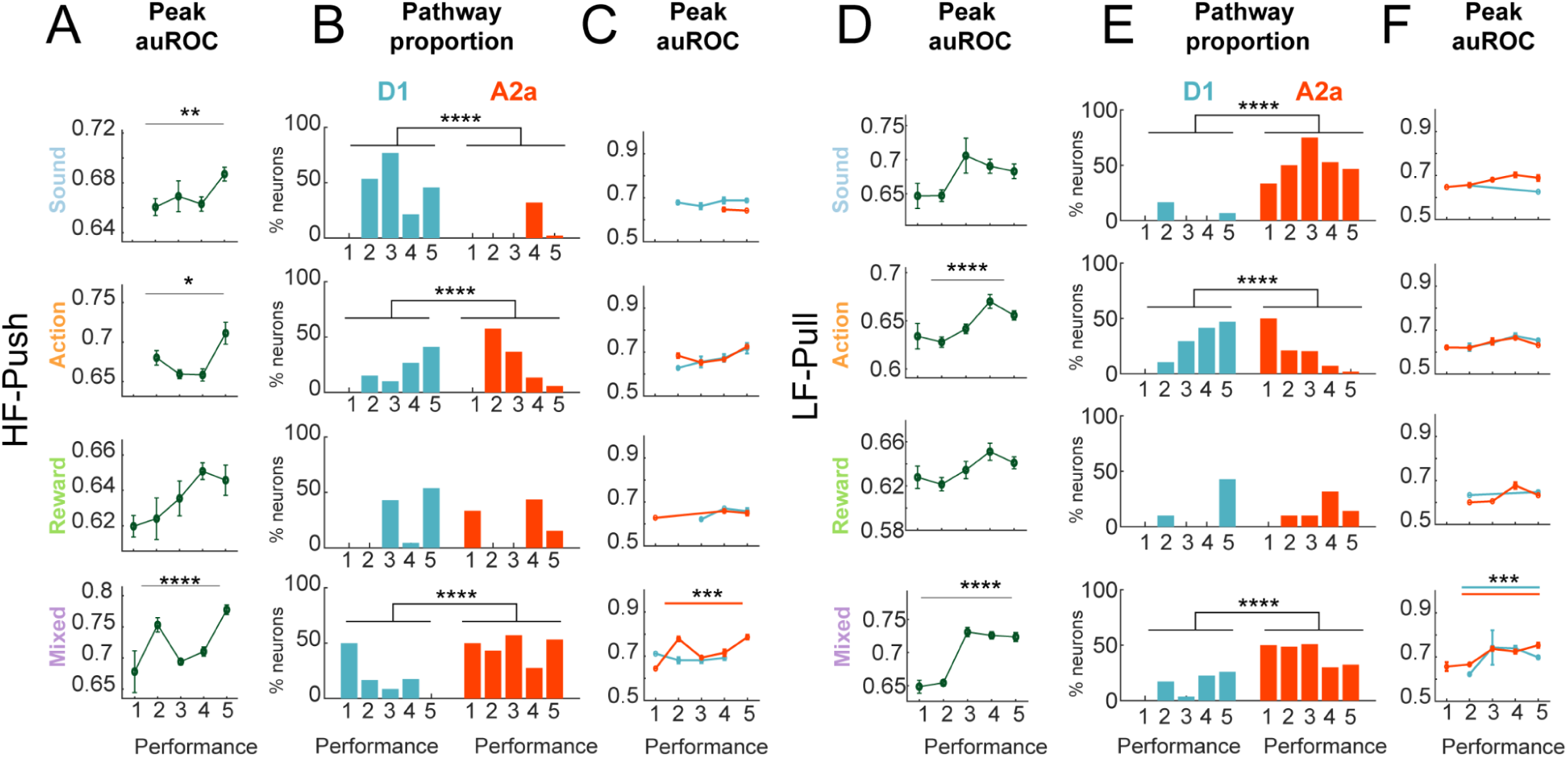
Category tuning strengthens with learning and pathway routing is category and association dependent. **A**, HF-Push association, peak tuning (maximum absolute auROC) across five representative training days, showing learned related increases in peak tuning for sound, action and mixed neurons(Kruskal-Wallis test across days, p<0.05). **B**, Hf-Push association, pathway composition of category neurons across learning, restricting to neurons with confirmed pathway identity(D1 vs A2a). Sound neurons are enriched in D1, action neurons become D1 biased with training, and mixed neurons predominantly A2a. **C**, HF-Push association, peak tuning across learning split by pathways, showing a selective learning related increase in peak tuning for a2a neurons. **D,** LF-Pull association. Peak tuning across training days, showing learning-related increases in peak tuning for Action and Mixed neurons (Kruskal-Wallis test across days, p < 0.05). **E,** LF-Pull association. Pathway composition of category neurons across learning (confirmed D1 vs A2a). Action neurons show a robust shift toward D1 with training, whereas Mixed neurons are predominantly A2a; Sound neurons show only a weak and non-significant trend due to low counts (Fisher-Monte Carlo; see Results). **F,** LF-Pull association. Peak tuning across learning split by pathway, showing that Mixed neurons in both pathways increase tuning with training.

We next examined how these category signals were distributed across pathways. Restricting to neurons with confirmed identity (D1 vs A2a), we compared pathway composition within each functional category across learning (Fig. 4B,E). In the Sound category, HF-Push was strongly D1-enriched (45 D1 vs 10 A2a; exact binomial vs 50%, p < 0.001). In LF-Pull, Sound neurons were instead strongly A2a-enriched overall (2 D1 vs 24 A2a; exact binomial vs 50%, p < 0.001), although their distribution across days did not vary significantly (Fisher-Monte Carlo, p = 0.63). In contrast, Action neurons showed robust pathway differences in both associations (Fisher-Monte Carlo, p ≤ 0.001), with the fraction of D1 cells increasing across training. Mixed neurons were strongly A2a-biased overall (HF-Push: 18 D1 vs 99 A2a; LF-Pull: 46 D1 vs 96 A2a; both p < 0.001, exact binomial vs 50%). Reward neurons showed smaller and less consistent pathway differences.

Finally, we tested whether learning-related changes in peak tuning differed by pathway within each category. In HF-Push, the clearest tuning increase with learning occurred in A2a Mixed neurons (Fig. 4C, Kruskal Wallis, p < 0.001). In LF-Pull, Mixed neurons in both pathways increased tuning across learning (Fig. 4F, Kruskal Wallis, p < 0.001).

We also examined whether functional category neurons showed spatial clustering or systematic changes in modulation polarity with learning. Category neurons showed no reliable distance-similarity structure in either association, and distance-similarity coupling did not change across learning (Supplementary Fig. 3A,B,E,F). In contrast, the balance of up- versus down-modulated neurons shifted with performance in a category- and association-dependent manner, with limited pathway-specific differences (Supplementary Fig. 3C,D,G,H).

In summary, learning sharpened category-specific modulation and re-routed signals across pathways in an association-dependent manner. Peak tuning (|auROC|) increased with performance for Sound/Action/Mixed in HF-Push and for Action/Mixed in LF-Pull, and the Action category showed more up-modulated neurons at higher performance in both associations. Pathway composition dissociated by function: Sound was D1-enriched in HF-Push but A2a-enriched in LF-Pull; Action shifted progressively toward D1 with training in both associations; and Mixed remained A2a-biased overall, with stronger tuning in A2a neurons. Reward category showed no consistent differences, and functional categories were spatially intermingled rather than segregated into local clusters. Together, these results suggest that TS SPN population codes evolve with learning by strengthening task-tuned activity and, in parallel, shifting which pathway carries specific task-related information.

To move beyond epoch-averaged modulation and ask which trial-by-trial variables best explain TS responses, we fit generalized linear models (GLMs) to trial-wise peak ΔF/F (change in fluorescence relative to baseline fluorescence) within three task-aligned windows: Sound, Action, and Outcome (see Methods; model validation in Supplementary Fig. 4). For each neuron, day, and association, peak responses were modeled using predictors for current-trial Action (movement vs omission), Reward (delivered vs not), and within-session trends (TrialIdx), with optional first-order history terms (PrevAction, PrevReward).

In the Action window, GLM coefficients shifted with learning in both associations. In LF–Pull, the reward coefficient (βReward) increased while the action coefficient (βAction) decreased across sessions (Fig. 5A; one-way ANOVA across sessions, p < 0.05), such that by late training peak responses were more strongly associated with reward delivery than with movement versus omission. The same learning-dependent pattern was present in HF–Push (Fig. 5F; one-way ANOVA across sessions, p < 0.05). Example neurons illustrate these effects: one neuron showed larger peak responses on rewarded versus unrewarded action trials at high performance (LF-Pull: Fig. 5B; HF-Push: Fig. 5G), whereas another showed strong differences between action trials and omission trials early in learning that diminished at high performance (LF-Pull: Fig. 5C; HF-Push: Fig. 5H).

**Figure 5.**
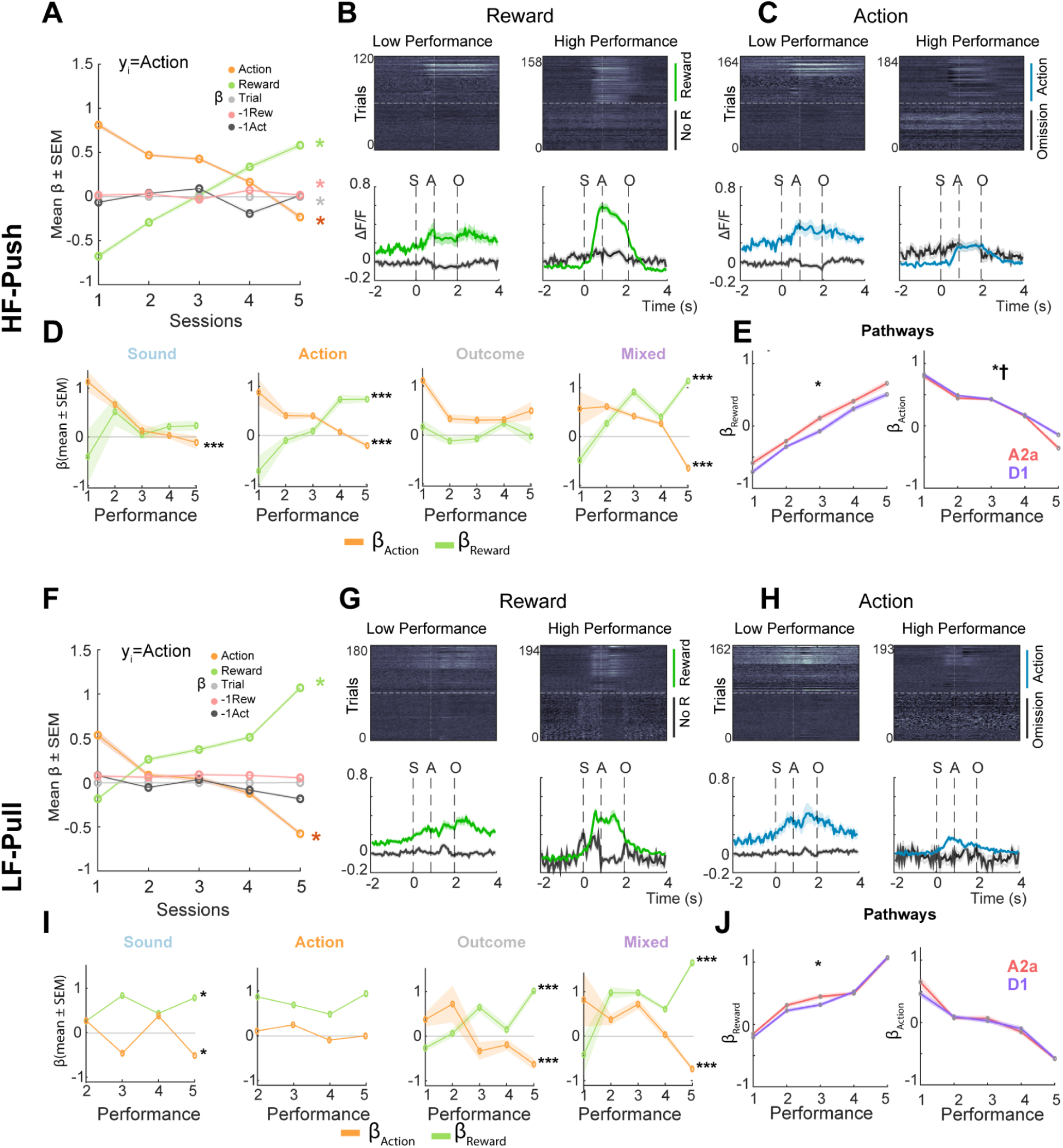
Learning shifts Action-window predictors from movement-related to reward-related contributions. **A,** HF-Push. GLM β coefficients for predicting trial-wise peak ΔF/F in the Action window across five representative sessions, showing a learning-dependent increase in βReward and decrease in βAction (see Results; one-way ANOVA across sessions). **B,** HF-Push. Example neuron illustrating stronger peak responses on rewarded versus unrewarded action trials at high performance. **C,** HF–Push. Example neuron illustrating strong differences between action trials and omission trials early in learning that diminish at high performance. **D,** HF-Push. βReward and βAction grouped by functional category (Sound, Action, Reward, Mixed), highlighting the category subsets that most clearly reflect the learning-dependent coefficient shift (see Results; one-way ANOVA across sessions). **E,** HF-Push. Pathway split (D1 vs A2a) for Reward β (left) and Action β (right) across sessions. Two-way ANOVA (Pathway × Session; categorical) shows pathway effects for both coefficients, with pathway-specific learning dynamics evident for Action β (see Results). **F,** LF-Pull. Same as A for the LF-Pull association, showing a learning-dependent increase in Reward β and decrease in Action β (see Results; one-way ANOVA across sessions). **G,** LF-Pull. Example neuron illustrating stronger peak responses on rewarded vs unrewarded trials at high performance. **H,** LF-Pull. Example neuron illustrating strong movement vs omission modulation early in learning that diminishes at high performance. **I,** LF-Pull. Reward β and Action β grouped by functional category, identifying the category subsets that best recapitulate the learning-dependent coefficient shift (see Results; one-way ANOVA across sessions). **J,** LF-Pull. Pathway split (D1 vs A2a) for Reward β (left) and Action β (right) across sessions. Two-way ANOVA indicates a modest pathway offset for Reward β without evidence for pathway-specific learning dynamics across sessions, and no reliable pathway effects for Action β (see Results).

To identify which neurons contributed most to the population-level coefficient shifts observed in Fig. 5A,F, we grouped Action-window coefficients by functional category. In HF-Push, the learning-dependent increase in βReward and decrease in βAction was most prominent in Action and Mixed neurons (Fig. 5D; one-way ANOVA across sessions, p < 0.05). In LF-Pull, the same coefficient shift was strongest in Reward and Mixed neurons (Fig. 5I; one-way ANOVA across sessions, p < 0.05).

We next tested whether pathway differences (D1 vs A2a) were present and whether they changed across learning by running a two-way ANOVA on neuron-level Action-window coefficients with factors Pathway and Session (five sessions; Fig. 5E,J). In HF-Push, βReward differed by pathway (Pathway main effect p = 8.42 × 10⁻⁸) without a Pathway × Session interaction (p = 0.59477), whereas βAction showed both a pathway main effect (p = 0.001156) and a Pathway × Session interaction (p = 0.001431), indicating pathway-specific learning dynamics in the movement-related coefficient (Fig. 5E). In LF-Pull, βReward showed a modest overall pathway offset (Pathway main effect p = 0.03299) without evidence that the A2a–D1 difference changed across sessions (Pathway × Session p = 0.11707), and βAction showed no pathway main effect (p = 0.26361) and no interaction (p = 0.09137) (Fig. 5J).

Model validation showed that predictive structure was concentrated in the Action window. Cross-validated R² (cvR²) in the Sound and Outcome windows stayed near the null bands (Supplementary Fig. 4E,G), whereas Task-only cvR² values in the Action window lay clearly above both null baselines across days and associations (Supplementary Fig. 4F). Including history terms did not improve performance: compared with the Task-only model, the Full model (adding PrevAction and PrevReward) produced slightly lower cvR² on average (Supplementary Fig. 4A-D), and unique contribution analysis showed near-zero contributions from PrevAction and PrevReward (Supplementary Fig. 4E-G). Instead, Reward and Action accounted for most of the explained variance in the Action window, with TrialIdx contributing more modestly.

Taken together, the GLM analysis adds a trial-by-trial view of TS encoding that complements the auROC/category framework: it shows that learning not only recruits more task-modulated neurons, but also reweights which task variables are most closely related to action-epoch responses, shifting from movement-related to reward-related influences as animals reach high performance.

In the stimulus-response (go/omit) task, associations were trained in separate sessions, allowing us to track how TS activity is recruited across sound, action, and reward epochs as animals learn a stable sound-action association. However, because the tone-action assignment is constant within a session and only one association is trained at a time, this design does not test within-session selection between opposing actions based on sound identity. We therefore trained a separate cohort on a two-choice (2AFC) task, in which both tones were interleaved within the same session and mice selected between push and pull on every trial based on sound identity (Fig. 6A; recording sites, Supplementary Fig. 5A).

**Figure 6.**
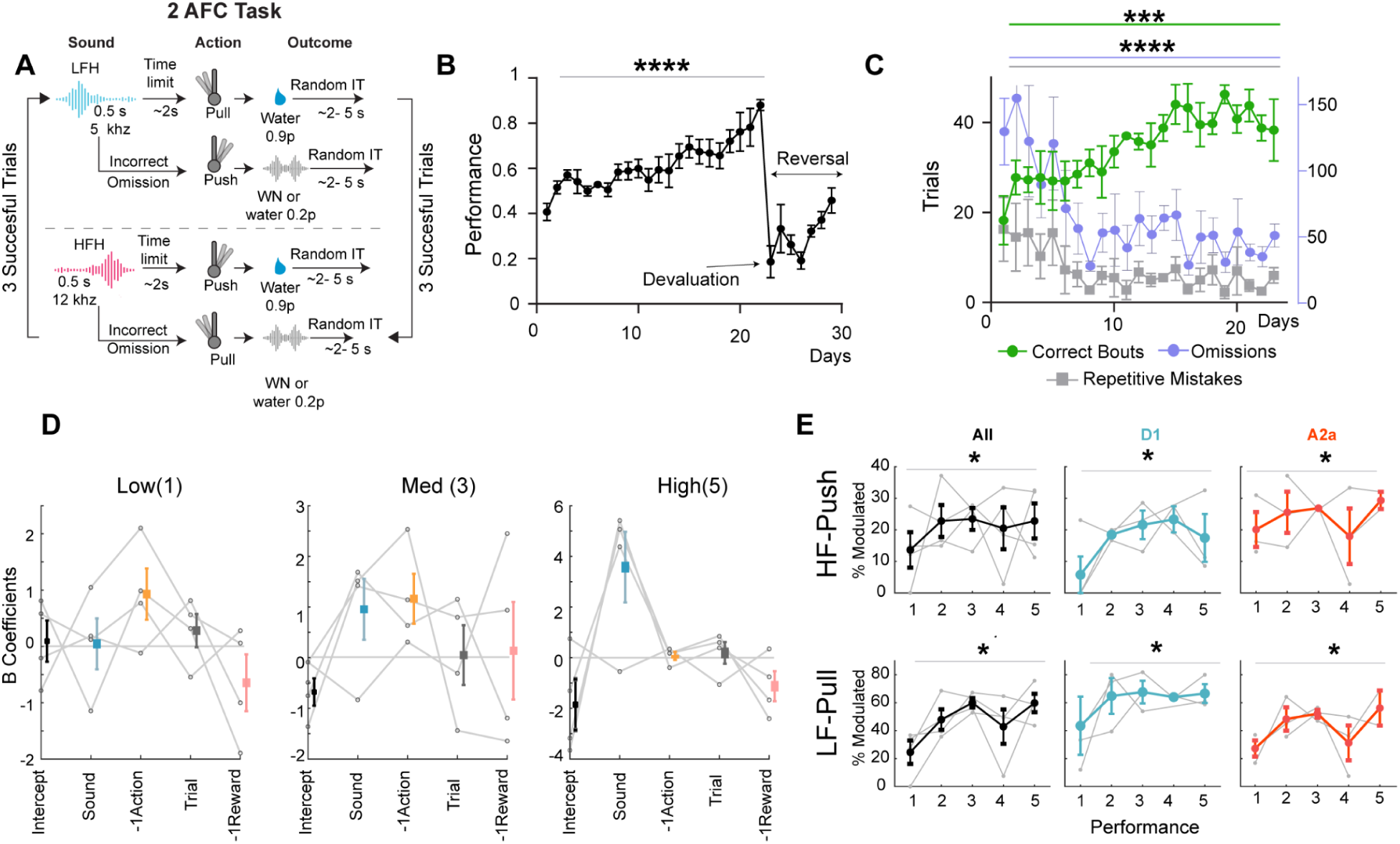
| Mice learn 2AFC task and TS modulation increases with performance. **A,** 2AFC task. Both tones were interleaved within the same session, and mice selected between push and pull on every trial based on sound identity. LF cued pull and HF cued push; in a separate reversal session, the assignment was switched. Correct responses delivered water with 0.9 probability, whereas incorrect responses triggered white noise and the trial restarted. **B,** Behavioral performance (Correct / (Correct + Incorrect)) increased across training days and dropped in the reversal session (Kruskal-Wallis across days, p < 0.001). **C,** Learning metrics across training. Correct bouts (runs of ≥2 correct trials) increased (Kruskal-Wallis, p < 0.05), while repetitive errors (≥2 consecutive incorrect trials) and omissions decreased (Kruskal-Wallis, p < 0.001). **D,** Behavioral GLM coefficients across performance levels, showing an increase in the β coefficient for current sound on high-performance days. **E,** TS neural modulation across five performance days in the 2AFC task. The fraction of auROC-defined task-modulated neurons increased with performance (Kruskal-Wallis, p < 0.05), with no reliable difference between D1 and A2a in the overall modulation metric.

In the 2AFC task, the tone-action assignment was LF-Pull and HF-Push, with both trial types presented within the same session; in a separate reversal session, the assignment was switched to LF-Push and HF-Pull. Correct responses delivered water with 0.9 probability (0.1 probability of no reward), whereas incorrect responses triggered white noise and the trial restarted. The inter-trial interval was 2-5 s, the response window was 5 s, and the tone lasted 500 ms (Fig. 6A).

Behavioral performance, defined as Correct / (Correct + Incorrect), increased across training days (Kruskal-Wallis, p < 0.001) and dropped in the reversal session (LF-Push, HF-Pull; Fig. 6B). To further quantify learning, we measured correct bouts (runs of ≥2 correct trials), repetitive errors (≥2 consecutive incorrect trials), and omissions. Correct bouts increased (Kruskal-Wallis, p < 0.05; Fig. 6C), while repetitive errors and omissions declined with training (Kruskal-Wallis, p < 0.001; Fig. 6C).

To probe decision strategy, we summarized behavior at low, medium, and high performance (performance ≈ 0.4, 0.6, and 0.8, respectively) and fit a behavioral GLM with predictors for current sound, previous action, trial index, and previous reward. On high-performance days, the β coefficient for current sound increased, consistent with choices being driven primarily by current auditory evidence rather than recent history (Fig. 6D).

Across the 2AFC task, we selected five representative sessions ranked from lowest to highest performance to assess TS neural modulation using auROC relative to baseline within the behavioral windows. Mean performance values (mean ± s.e.m.) across Sessions 1–5 were 0.44 ± 0.03, 0.62 ± 0.05, 0.68 ± 0.06, 0.78 ± 0.06, and 0.85 ± 0.05, respectively. As in the go/omit task, the fraction of modulated neurons increased with performance in both pathways (Fig. 6E; Kruskal-Wallis, p < 0.05).

We tested whether individual TS SPNs distinguish between the two sound-action associations within the same task. We computed an association-selectivity auROC by comparing correct trials from the LF-Pull versus HF-Push associations in two time windows: one locked to the sound epoch and one spanning the action/reward epoch. Neurons with auROC values significantly above chance (vs shuffle) were labeled LF-Pull preferring, those significantly below chance were labeled HF-Push preferring, and neurons that did not significantly discriminate between associations, but were modulated, were labeled non-association modulated (Fig. 7A; Supplementary Fig. 5B).

**Figure 7.**
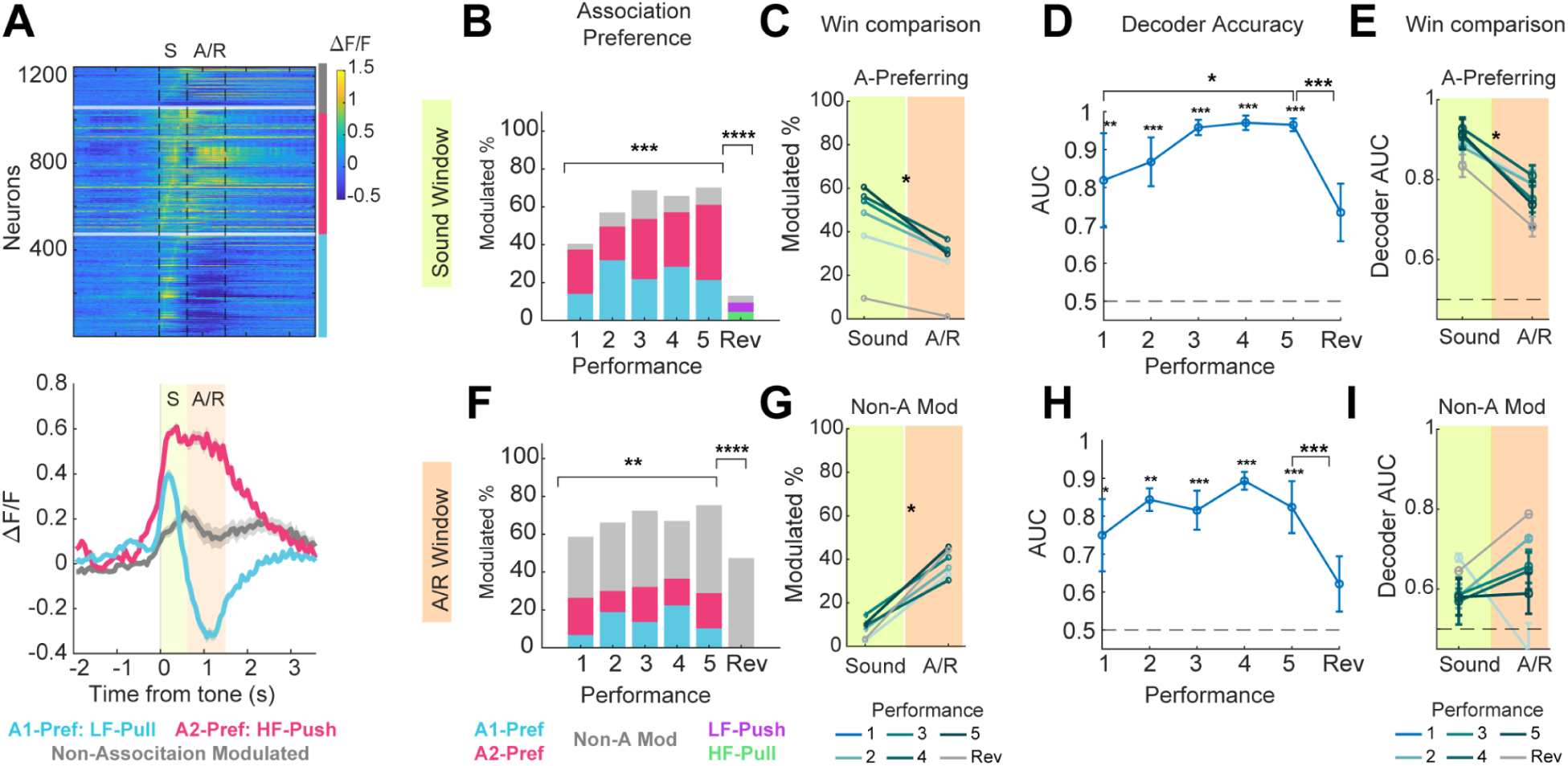
Learning increases sound-window association-preferring neurons and decoding accuracy. **A,** Association selectivity in the 2AFC task. TS SPNs were classified as A1-Pref., A2-Pref., or Non-A Mod. using an association-selectivity auROC comparing correct LF-Pull (A1) versus HF-Push (A2) trials in sound-locked and action/reward windows (significance assessed against shuffle). **B,** Sound window. The fraction of association-preferring neurons increased across training days and decreased after reversal (χ², p < 0.001). **C,** Window comparison of association-preferring fractions. Sound-locked responses contained a higher fraction of association-preferring neurons than the action/reward window (paired Wilcoxon signed-rank, p < 0.05). **D,** Sound-window decoding. Decoder performance exceeded a label-shuffle null on most days (1,000 shuffles; p < 0.05), improved across training (linear mixed-effects model; see Results), and declined after reversal (paired Wilcoxon signed-rank, p < 0.001). **E,** Decoding comparison for association-preferring neurons. Decoding was higher in the sound window than in the action/reward window (paired Wilcoxon signed-rank, p < 0.05). **F,** Action/reward window. Association-preferring neurons were present, changed across training (χ², p < 0.01), and decreased after reversal (χ², p < 0.001). **G,** Window comparison of Non-A Mod. fractions. Non-A Mod. neurons were more prevalent in the action/reward window than in the sound window (paired Wilcoxon signed-rank, p < 0.05). **H,** Action/reward decoding. Decoder performance exceeded the shuffle-null on most days (1,000 shuffles; p < 0.05) but did not show a reliable increase across training (linear mixed-effects model; see Results). **I,** Decoding using Non-A Mod. neurons. Decoder AUC did not differ reliably between the sound and action/reward windows (paired Wilcoxon signed-rank; n.s.).

In the sound window, the fraction of association-preferring neurons increased across training days and decreased after reversal (χ², p < 0.001 for both; Fig. 7B). Consistent with this, sound-locked responses contained a higher fraction of association-preferring neurons than the action/reward window (paired Wilcoxon signed-rank, p < 0.05; Fig. 7C). An L2-regularized logistic regression classifier trained within each session to distinguish LF–Pull versus HF–Push trials from sound-locked population responses mirrored these effects: decoding exceeded a label-shuffle null on most days (p < 0.05; 1,000 shuffles; Fig. 7D; Supplementary Fig. 5C,D), improved across training (linear mixed-effects model; slope = 0.034 AUC/day, p = 0.0287), and declined after reversal (paired Wilcoxon signed-rank, p < 0.001; Fig. 7D). Decoding within association-preferring neurons was correspondingly higher in the sound window than in the action/reward window (paired Wilcoxon signed-rank, p < 0.05; Fig. 7E).

In the action/reward window, association-preferring neurons were also present and changed across training (χ², p < 0.01), and their fraction decreased after reversal (χ², p < 0.001; Fig. 7F). Non-association modulated neurons were more prevalent in the action/reward window than in the sound window (paired Wilcoxon signed-rank, p < 0.05; Fig. 7G). Decoding in the action/reward window exceeded the shuffle-null on most days (p < 0.05; 1,000 shuffles; Fig. 7H; Supplementary Fig. 5E,F) but did not show a reliable increase across training (linear mixed-effects model; slope = 0.0198 AUC/day, p = 0.253). In contrast to association-preferring neurons, decoding using non-association modulated neurons was weak and did not differ reliably between the sound and action/reward windows (Fig. 7I).

Finally, we asked whether association preference differed by pathway. Pathway proportions (D1 vs A2a) were similar across association-preferring and non–association-modulated groups in both windows (paired Wilcoxon signed-rank, p > 0.05; Supplementary Fig. 5G). A small window-by-pathway difference was observed for HF-Push-preferring neurons, with more A2a in the sound window and more D1 in the action/reward window (Wilcoxon signed-rank, p < 0.05; Supplementary Fig. 5H).

Together, these results show that TS carries robust association information during the two-choice task, with the sound window providing the strongest and most learning-dependent signal: association-preferring fractions and sound-window decoding increase with training and drop after reversal, whereas action/reward decoding is present but less performance-dependent.

A key open question is whether TS representations are organized in space and stable over time. To address this, we combined our spatial and cross-day analyses across the stimulus-response (go/omit) task and the two-choice (2AFC) task.

In the go/omit task, functional category neurons did not show evidence of spatial clustering. Pairwise functional similarity did not reliably relate to intersomatic distance in either association (HF-Push and LF-Pull; Fig. 8A,B; Supplementary Fig. 3; all p > 0.05), indicating that category coding is spatially intermingled at this scale.

**Figure 8.**
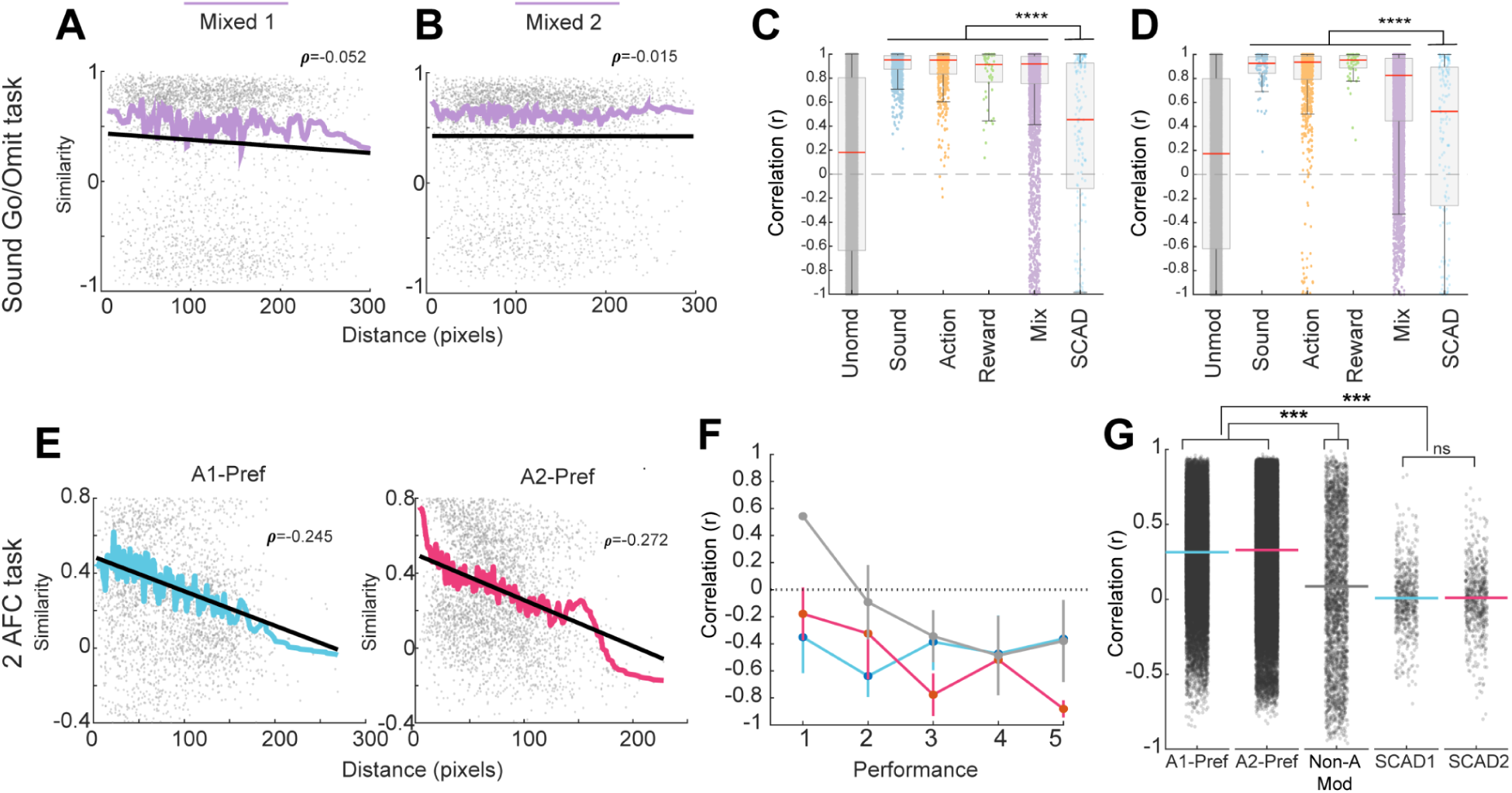
Stable population-level structure with drifting single-cell participation, and local spatial organization of association-preferring activity. **A,** Go/omit task (HF-Push), Mixed category. Pairwise functional similarity as a function of intersomatic distance, showing a weak negative relationship (Pearson r = −0.052, p < 0.01, slope = −0.0006). The solid colored curve shows a LOWESS smoother, and the black line shows the linear fit. **B,** Go/omit task (LF-Pull), Mixed category. Same as A (Pearson r = −0.015, p = 0.347, slope = 0). **C,** HF-Push cross-day stability Go/Omit task. Cross-day pairwise correlations of response fingerprints for neurons grouped by functional category compared with correlations for SCAD-matched same neurons across days and unmodulated neuron pairs. Within-category similarity remained high (medians: Sound 0.95, Action 0.95, Reward 0.92, Mixed 0.92), SCAD correlations were intermediate (median 0.45), and unmodulated pairs clustered near baseline (median 0.18). **D,** LF-Pull cross-day stability Go/Omit task. Same as C for LF-Pull (medians: Sound 0.93, Action 0.94, Reward 0.95, Mixed 0.83; SCAD median 0.52; unmodulated median 0.17). **E,** 2AFC task (sound window). Spatial organization of association-preferring neurons. Pairwise functional similarity decreased with intersomatic distance for both A1-Pref. neurons (A1 = LF-Pull; Pearson r = −0.245, p < 0.001, slope = −0.478) and A2-Pref. neurons (A2 = HF–Push; Pearson r = −0.272, p < 0.001, slope = −0.489). The solid colored curve shows a LOWESS smoother, and the black line shows the linear fit. **F,** Distance-similarity relationship across performance days in the 2AFC task, remaining negative across learning for both association-preferring populations. **G,** Cross-day association structure versus same-cell stability. Cross-day similarity (ρ) was higher for association-preferring neurons than for Non-A Mod. neurons (A1-Pref. vs Non-A Mod.: Δmedian = 0.232, BH-FDR-adjusted p < 0.001; A2-Pref. vs Non-A Mod.: Δmedian = 0.265, BH-FDR-adjusted p < 0.001) and exceeded SCAD controls (all BH-FDR-adjusted p < 0.001), whereas SCAD control groups did not differ (BH-FDR-adjusted p = 0.66).

We next asked whether category-level response structure persists across days. Using same cells across days (SCAD) tracking (matched ROIs across sessions), we obtained ∼40-60% cell retention across the five representative performance days in both associations (Supplementary Fig. 2G-J). Within-category similarity remained high across days, whereas SCAD-matched same-cell correlations were intermediate and unmodulated pairs clustered near baseline (Fig. 8C,D). Thus, category-level output structure is stable across learning even though single-neuron response patterns drift across days.

We then asked whether the same population versus single-cell pattern holds for association representations in the 2AFC task. Association-preferring neurons in the sound window showed clear spatial structure: pairwise similarity decreased with distance for both LF-Pull preferring and HF-Push preferring neurons (Fig. 8E; Pearson correlation, p < 0.001), and this negative distance similarity relationship was maintained across performance days (Fig. 8F).

Finally, we quantified cross-day association structure by computing cross-day similarity (ρ) from correlations of ΔF/F response time courses across day pairs. Association-preferring neurons showed higher cross-day similarity than non-association modulated neurons and exceeded SCAD controls (p < 0.001), whereas SCAD control groups did not differ (Fig. 8G; BH-FDR-adjusted comparisons). Together, these analyses highlight a shared principle across tasks stable population-level structure with more variable single-cell participation—while also revealing a dissociation in spatial organization: category coding is distributed, whereas association-preferring activity is locally structured in space.

## Discussion

We set out to understand how learning transforms initially meaningless sounds into cues that guide action. By tracking pathway-identified TS SPNs longitudinally, we show that learning recruits more TS neurons across sound, action, and outcome epochs, strengthens tuning, and builds robust association information when animals must flexibly apply sound-action association. At the same time, functional category structure remains stable across days even as the identities of responsive neurons drift. Learning also routes signals differently across pathways in an association and epoch dependent manner, with action-related signals increasingly concentrated in D1 SPNs and multi-epoch Mixed signals biased toward A2a SPNs. Together, these findings support TS as a learning-related hub in which sound-action behavior is implemented through coordinated recruitment across trial epochs and pathway reweighting.

### Learning recruits TS beyond a global motor command

As training progressed, a larger fraction of TS neurons became task-modulated and reliably distinguished correct trials from omissions, with stronger activity when animals were engaged and executed the required action. Importantly, TS population activity was not dominated by a single, global motor command: movement onset alone showed no reliable population-wide coupling (Supplementary Fig. 1C), consistent with TS reflecting task structure beyond movement initiation. A plausible account is that learning progressively increases the gain of TS circuits as cues acquire behavioral meaning, via convergent sensory drive and neuromodulatory signals that support cue salience and engagement (Menegas et al., 2018; Green et al., 2024). This view fits with evidence that posterior/auditory striatum rapidly develops sound-related codes during associative learning (Znamenskiy and Zador, 2013; Guo et al., 2018), and with TS microcircuit mechanisms that can calibrate recruitment during sound processing through inhibition (Li et al., 2024; Druart et al., 2024).

### Stable category structure with drifting single-cell identities

A central result is that TS maintains stable category-level organization across learning (Sound/Action/Outcome/Mixed), even as individually tracked cells show weaker cross-day consistency. Similar representational drift has been reported in sensory cortex, whereas some motor circuits show stronger single-neuron stability, suggesting that stability-flexibility balances may be region and task dependent (Deitch et al., 2021; Bauer et al., 2024; Jensen et al., 2022). Our results fit models in which networks maintain stable outputs while internal activity patterns evolve, rather than relying on fixed single-cell templates (Maass et al., 2002; Driscoll et al., 2022). In this view, downstream structures may read out a stable TS code even as individual neurons rotate through it over learning and changing behavioral state.

### Learning sharpens tuning and routes information across pathways

Learning strengthened category tuning in an epoch and association dependent way, and pathway routing dissociated by function. The progressive enrichment of D1 neurons in the Action category aligns with basal ganglia frameworks in which the direct pathway supports selection and execution of the chosen action, while the indirect pathway contributes to suppression and control of alternatives (Albin et al., 1989; DeLong, 1990; Kravitz et al., 2010; Mink, 1996; Yttri & Dudman, 2016; Tecuapetla et al., 2016; Cruz et al., 2022). The A2a bias among Mixed neurons suggests that indirect-pathway populations carry multi-epoch information spanning sensory, action, and outcome structure, consistent with roles in stabilizing selection, inhibitory control, and context appropriate suppression (Mink, 1996; Yttri & Dudman, 2016; Cruz et al., 2022).

At the same time, sound-related pathway routing depended on the learned association: Sound neurons were D1-enriched in HF-Push but not in LF-Pull. This argues against a fixed “sensory-to one pathway” channel and instead supports flexible routing shaped by the learned contingency and local circuit context. TS receives complementary auditory inputs from cortex and thalamus (Ponvert & Jaramillo, 2019; Chen et al., 2019), and recent physiology shows a subthreshold bias favoring D1 SPNs that can arise from pathway-specific inhibitory gating (Druart et al., 2024). PV interneurons provide another lever over auditory responses and auditory-guided behavior in TS (Li et al., 2024), and could help shape association-dependent recruitment. Notably, even though TS exhibits sensory topography and frequency selectivity (Guo et al., 2018; Chen et al., 2019), TS activity can still differentiate contingencies, indicating that association information is embedded beyond pure sensory gradients.

The learning-related increase in down-modulated Sound neurons further supports a role for inhibitory gating in TS learning, consistent with PV interneurons shaping auditory responses and with pathway-specific excitatory/inhibitory balance (Li et al., 2024; Druart et al., 2024). Finally, category neurons were spatially intermingled rather than clustered, consistent with distributed category coding across the imaged fields. This contrasts with the spatial organization of association-preferring neurons, suggesting that category structure and association structure may be implemented at different spatial scales.

### Trial-wise modeling reveals learning-dependent reweighting during action execution

Using GLMs on trial-wise peak ΔF/F, predictive structure was strongest in the Action window; task variables predicted action window peaks above null baselines, whereas Sound and Outcome windows hovered near null when using peak-based summaries. Across learning, the Reward term gained predictive weight while the Action term diminished, consistent with a shift from movement-driven to reward-driven modulation as performance improved. This complements the broader view that striatal activity integrates action and outcome information as learned behavior stabilizes, and is consistent with dopamine-linked valuation/salience signals in TS (Menegas et al., 2018; Green et al., 2024; Gerfen and Surmeier, 2011; Balleine and O’Doherty, 2010). Our pathway by session analysis further indicates that, in general, most pathway effects reflect stable offsets rather than diverging trajectories.

History terms (previous action and reward) contributed little unique variance and slightly reduced generalization. We interpret this cautiously as a limitation of peak-based metrics rather than evidence that history is irrelevant, because history dependence may reside in timing, sustained dynamics, or latent states such as engagement or arousal that are not captured by peak ΔF/F, and may vary across striatal subregions and task demands (Valjent and Gangarossa, 2021; Menegas et al., 2018).

### Two-choice task: sound-locked TS activity carries the strongest association information

When animals performed the two-choice (2AFC) task, behavior improved with training and choices became more strongly driven by current auditory evidence, consistent with increased reliance on the sensory cue rather than recent history. TS SPNs carried robust association information, but this information was strongest and most learning-dependent in the sound window; learning increased the fraction of sound-window association-preferring neurons and improved sound-window decoding, both of which dropped after reversal. Association-related signals were also present in the action/reward window, but decoding there did not reliably increase with training, and non-association modulated neurons were more prevalent than in the sound window. Together, these results argue that TS reflects the learned association between sound and action, with the sensory phase carrying the clearest association information, consistent with prior demonstrations of stable sound codes supporting decisions in posterior/auditory striatum (Guo et al., 2018) and with emerging pathway-specific choice signals in auditory tasks (Tang et al., 2025).

### Association-preferring neurons show local structure despite single-cell drift

A striking dissociation emerges between category coding and association coding. Category neurons did not show a distance-similarity relationship, but sound-window association-preferring neurons did, with nearby neurons showing more similar responses than distant ones across learning. Such spatial structure aligns with prior work showing organized striatal population activity during behavior (Barbera et al., 2016; Klaus et al., 2017; Parker et al., 2018). Cross-day analyses further show that association-related population structure is more stable than the trial-by-trial patterns of the same individually tracked cells, consistent with representational drift at the single-cell level despite preserved association organization at the population level. This points to a TS code that is reliably readable at the population level while allowing cell-level reassignment over days.

### Circuit-level interpretation

Several circuit mechanisms could support progressive recruitment, pathway-specific routing, and stable population outputs. TS receives convergent auditory drive from cortex and thalamus with complementary coding (Ponvert and Jaramillo, 2019; Chen et al., 2019), and local inhibition can gate how those inputs are expressed in SPNs. PV interneurons in TS strongly regulate auditory responses and auditory-guided behavior (Li et al., 2024), and pathway-specific inhibitory routing can bias subthreshold dynamics across D1 and A2a neurons (Druart et al., 2024). In parallel, salience-biased dopamine signaling in TS may amplify learning-relevant cues and stabilize the currently favored association (Menegas et al., 2018; Green et al., 2024), while also providing an action prediction error signal that can act as a value-free teaching signal to reinforce repeated sound-action associations (Greenstreet et al., 2025). Together, complementary sensory inputs, PV-mediated gating, and dopaminergic teaching signals offer a plausible substrate by which TS could support stable, association-sensitive sound-action codes while allowing the specific neurons carrying those codes to drift.

### Limitations and future directions

Several limitations are worth noting. Pathway assignment relies on confirmed tdTomato+ cells and putative opposite-pathway classification for GCaMP-only cells. Although this strategy has been used successfully here and in other studies (Sheng et al., 2019; Maltese et al., 2021), it could misclassify a minority of neurons. Future approaches that directly validate pathway identity, such as Cre-on/Cre-off strategies (Saunders and Sabatini, 2015) or dual-color indicators (Wu et al., 2025), will help refine these assignments. Cross-day ROI matching is conservative and may undercount true retention, though it supports stable longitudinal sampling at the population level. The GLM uses peak ΔF/F and may miss history-dependent structure expressed in timing or sustained dynamics. Finally, the head-fixed two-tone joystick task constrains generality; extending these analyses to richer sound spaces, more complex action sets, and causal manipulations of thalamic input, PV interneurons, and dopamine signals will be important for testing the mechanistic hypotheses suggested here.

In summary, TS supports learning of sound-action associations by recruiting and sharpening task-tuned activity across epochs, routing information across pathways in an association-dependent manner, and building robust sensory-phase association coding that supports flexible behavior, all while maintaining stable population-level structure through drifting single-cell identities.

## Materials and Methods

### Experimental overview

We used longitudinal two-photon calcium imaging through GRIN lenses to track tail-of-the-striatum (TS) spiny projection neurons (SPNs) as head-fixed mice learned tone-guided joystick actions, and in a separate cohort, as they performed a two-choice (2AFC) sound-action task. We focused on three task-aligned epochs (Sound, Action, and reward) and quantified learning-related changes in task modulation, functional category structure (Sound/Action/Reward/Mixed), pathway routing (D1 vs A2a), and stability across days (including SCAD cell tracking). For the 2AFC cohort, we further quantified association selectivity, association decoding, and the spatial and cross-day structure of association representations.

### Animals

For single-association two-photon recordings, we used five double-transgenic mice (three A2a-Cre × Ai14: 2 male, 1 female; two D1-Cre × Ai14: 1 male, 1 female), in which tdTomato (Ai14) labels the Cre-defined pathway. For two-choice (2AFC) recordings, we used a separate cohort of four double-transgenic mice (two D1-Cre × Ai14: 1 male, 1 female; two A2a-Cre × Ai14: 2 male). Mice were maintained on a reverse light-dark cycle, socially housed with enrichment, and were ≥60 days old at the start of experiments. All procedures were approved by the Rutgers IACUC.

### Viral expression and surgical procedures

For the single-association cohort, we expressed GCaMP8f (Syn promoter); for the 2AFC cohort we expressed GCaMP7f (Syn promoter). Virus was diluted 1:1 in sterile saline. To distribute expression within TS, we delivered a total of ∼600 nL over ∼10 min, with an additional ∼10 min dwell time per injection, using six ∼100 nL injections spanning the intended imaging site. Stereotaxic targeting was centered at AP −1.2 mm and ML 2.6 mm (left hemisphere), with depths of 1.9-2.0 mm from the pial surface. The GCaMP variant differed between cohorts due to availability at the time of data collection; analyses were performed within cohorts.

A 1-mm GRIN lens was implanted over the injection site by enlarging the craniotomy, aspirating overlying tissue to accommodate the lens, lowering the lens to ∼1.9 mm (just above the transfected volume), and securing it with dental acrylic to form a protective cap. A custom 3D-printed guard protected the lens. Animals received local bupivacaine and standard postoperative analgesia, with daily health checks during recovery. At the end of experiments, we verified GRIN placement, viral expression, and implant location by standard histology and fluorescence imaging.

### Behavioral apparatus and joystick

Head-fixed mice were positioned in a restraining tube (Wagner et al., 2020) and interacted with a forelimb joystick mounted beneath the right paw. Joystick position was continuously recorded. Auditory stimuli were delivered from speakers positioned near the animal’s head. Water reward was delivered via a stainless-steel spout under solenoid-valve control. Task events, sound delivery, and reward timing were controlled by Arduino-based hardware and Python/MATLAB software. Hardware/software details for the open-source joystick platform are described in Linares-Garcia et al., 2025 and are available at: https://github.com/margolislab/Open-Source-Joystick-Platform

### Tasks, training, and behavioral measures

### Water restriction, habituation, and shaping

Mice were water-restricted and received water primarily during task sessions. Each reward delivered ∼0.01 µL of water; daily water intake was monitored and supplemented as needed to maintain body weight ≥80% of baseline. Habituation progressed from body-tube acclimation to gradual increases in head fixation up to ∼30 min per day. During a 3-5 day joystick-reward association phase, any joystick movement triggered ∼0.01 µL of water (up to ∼200 rewards per day).

### Single-association sound-action task

In the single-association task, tones were either low (5 kHz) or high (12 kHz), each presented with five overtones at 60 dB SPL for 500 ms. Trials began after a 2-5 s inter-trial interval (ITI) and included a 500 ms grace period followed by a 5 s response window. Sound-action associations were trained in separate sessions: low-frequency tones required a pull movement (LF-Pull) and high-frequency tones required a push movement (HF-Push). Correct responses yielded water with 0.9 probability (0.1 probability of white noise). Omission trials, in which no joystick displacement occurred within the response window, triggered 5 s of white noise before the next trial. Performance was quantified as Correct / (Correct + Omission).

### Two-choice (2AFC) task

In the 2AFC task, both tones were interleaved within the same session and mice selected between push and pull on every trial based on sound identity. Under the standard sound-action association, LF cued pull and HF cued push; in a separate reversal session, the association was switched to LF-Push and HF-Pull. Timing parameters were: inter-trial interval (2-5 s), tone duration (500 ms), and response window (5 s). Correct choices yielded water with 0.9 probability (0.1 probability of no reward). Incorrect choices triggered white noise and the trial restarted. Performance was calculated as Correct / (Correct + Incorrect). We summarized behavior using correct bouts (≥2 correct trials in a row), repetitive errors (≥2 incorrect trials in a row), and omissions.

To probe decision strategy in the 2AFC task, we modeled trial-by-trial choice with a logistic GLM including predictors for current sound (LF vs HF), previous action, trial index, and previous reward. Coefficients were estimated per animal and performance level (low/medium/high) and tested for changes across performance to assess whether choices became more sound-driven versus history-driven.

### Behavioral analyses

Behavioral performance metrics were computed offline from event logs and joystick traces. For the single-association task, performance was defined as Correct / (Correct + Omission); for the 2AFC task, as Correct / (Correct + Incorrect). Correct bouts were defined as runs of ≥2 correct trials, and repetitive errors as runs of ≥2 incorrect trials.

Joystick position was continuously sampled and low-pass filtered for analysis. Movement onset was detected when joystick displacement exceeded a fixed threshold above baseline. Reaction time was defined as the latency from tone onset to joystick movement onset. Reaction times were summarized per day and association and compared across days using Kruskal-Wallis tests.

### Two-photon imaging and acquisition

Imaging was performed with a Ti:sapphire Mai Tai DeepSee two-photon microscope tuned to 950 nm, enabling simultaneous excitation of GCaMP and visualization of tdTomato. Laser power at the sample was ∼1.5-1.8 mW. Images were acquired through the 1-mm GRIN lens with a 10× objective, yielding a field of view of roughly ±0.5 mm AP and ML, around the lens center. Mice were screened for robust expression and signal quality, and recordings commenced after up to ∼3 weeks of expression.

### Imaging preprocessing and ROI extraction

Movies were motion-corrected and segmented into ROIs with Suite2p. For each ROI, we used Suite2p’s local neuropil signal to correct somatic traces by subtracting a fixed fraction of neuropil from the raw ROI signal (contamination factor = 0.3). Analyses were restricted to ROIs classified as putative neurons by Suite2p’s cell-probability filter. Corrected traces were converted to ΔF/F relative to each ROI’s baseline fluorescence and subsequently z-scored for population analyses. ROIs that did not meet basic quality criteria (signal-to-noise, size, or stability) were excluded.

### Cross-day ROI tracking (SCAD)

To follow the same neurons across days, we generated mean reference images for each session and manually matched ROIs across sessions by centroid position and contour/shape similarity. Matched ROIs received a persistent unit identifier and were curated in a tracking table. This conservative, human-verified alignment minimized false matches while acknowledging small residual registration uncertainty.

### Task epoch definitions

We defined three task-aligned analysis windows for the single-association task: Sound (0-600 ms from tone onset; tone + grace period), Action (a fixed 1 s window time-normalized to movement onset by linear resampling for variable trial lengths), and reward (1,000 ms after reward delivery). Unless stated otherwise, baseline activity was computed from a pre-tone interval within the same trial.

### Pathway assignment

Pathway identity was confirmed by tdTomato expression in the appropriate Cre line (D1 in D1-Cre; A2a in A2a-Cre). When available, Suite2p’s red-channel probability was used with a ≥0.7 threshold to assign identity. For population summaries, GCaMP-only (“green”) cells were treated as putative opposite-pathway SPNs, whereas pathway-specific tests were restricted to confirmed red cells where indicated.

### Modulation, functional categories, and tuning auROC modulation and polarity

For each neuron and window, we computed an auROC value comparing within-window activity to a pre-tone baseline and assessed significance using 1,000 label permutations per neuron per window (α = 0.05). Polarity was defined as up-modulated (auROC > 0.6) or down-modulated (auROC < 0.4). Sliding-window analyses were used as a robustness check for the choice of fixed windows.

### Category assignment

Neurons were assigned to four non-overlapping functional categories based on which epoch windows showed significant modulation: Sound-only, Action-only, Outcome-only, or Mixed (significant modulation in more than one window).

### Peak tuning

For each neuron, we quantified peak tuning as the maximum absolute auROC value within its category window.

### Population and stability analyses (single-association) Motor coupling control

To assess motor coupling, we aligned population-mean neural activity to movement onset and computed the zero-lag Pearson correlation (r₀) and the full cross-correlogram between joystick magnitude and neural activity. Within-session significance was evaluated with circular-shift permutations that preserved slow drifts but disrupted trial timing, and group-level significance was assessed with Wilcoxon signed-rank tests across animals.

### Correct versus omission contrasts

Correct versus omission contrasts were computed as mean ΔF/F differences within modulated neurons for each epoch window.

### Category stability versus single-cell drift

To compare category stability with single-cell drift, we computed cross-day correlations of sign-aligned response fingerprints for neurons grouped by functional category (Sound, Action, Outcome, Mixed) and contrasted these with correlations for the same tracked cells across days (SCAD).

### Spatial organization of functional categories

To test for spatial organization, we computed pairwise functional similarity (Pearson correlation of trial-averaged responses within a window) for neuron pairs within a category and related it to intersomatic distance within the imaging field. Similarity-distance relationships were quantified with correlation coefficients and evaluated across performance days.

### GLM analyses (single-association)

For each neuron, day, and association, we modeled trial-wise peak ΔF/F within the Sound, Action, and Outcome windows using generalized linear models. Predictors were: Action (movement vs omission), Reward (delivered vs not), TrialIdx (within-session trial index), and two first-order history terms (PrevAction and PrevReward; set to zero on the first trial). We fit a “Full” model including all predictors and a “Task-only” model omitting the history terms.

Model quality was assessed with cross-validated R² (cvR²) using stratified folds within session and compared against two nulls: a circular-shift null that preserved slow drifts but broke trial timing, and a label-shuffle null that destroyed trial structure. Unique contributions were computed as the increase in cvR² when adding a given predictor to a model that already contained the remaining regressors, using identical cross-validation folds. Coefficients and unique contributions were summarized by functional category (Sound, Action, Outcome, Mixed) and pathway (D1, A2a).

### 2AFC analyses: association selectivity, decoding, and similarity

### Time windows and trial selection

All DRS analyses were run per session and per predefined time window. We defined a sound-locked window (0-600 ms from tone onset) and an action/reward window (600-1,500 ms from tone onset). Unless stated otherwise, analyses were restricted to correct trials and used trial-wise ΔF/F traces from each neuron.

### auROC-based association selectivity

To determine whether neurons preferred one sound-action association over the other, we computed, for each neuron and window, trial-wise ΔF/F responses on correct LF-Pull and HF-Push trials and calculated an auROC comparing the two trial distributions. By construction, auROC = 0.5 corresponds to chance; auROC > 0.5 indicates higher responses on LF-Pull than HF-Push trials, whereas auROC < 0.5 indicates the opposite.

Significance was assessed by permutation of association labels within session (10,000 shuffles). Neurons with auROC > 0.5 and *p* < 0.05 were classified as LF-Pull preferring; neurons with auROC < 0.5 and *p* < 0.05 were classified as HF-Push preferring. Neurons that did not significantly discriminate between associations but showed significant modulation versus baseline were classified as non-association modulated; the remainder were labeled unmodulated.

### Population association decoder

To quantify association information at the population level, we constructed for each trial a population vector comprising each neuron’s mean ΔF/F within the analysis window. We trained an L2-regularized logistic regression classifier to distinguish LF-Pull versus HF-Push trials using stratified k-fold cross-validation within session (k = 5), repeating cross-validation with new fold assignments. Within each fold, features were z-scored using training-set mean and standard deviation, and the same transform was applied to held-out trials. To avoid class-imbalance effects, trials were balanced by subsampling to an equal number per association. Decoder performance was quantified as ROC-AUC (0.5 = chance, 1.0 = perfect classification) on held-out trials and averaged across folds (and repeats).

To assess significance, we generated a label-shuffle null distribution by repeating the decoding procedure with permuted association labels within the session (typically 1,000 shuffles). We computed an empirical two-sided, chance-centered p-value as the fraction of shuffled AUCs whose deviation from chance (0.5) was at least as large as the observed deviation:

p = mean(|AUC_shuffle − 0.5| ≥ |AUC_obs − 0.5|).

### Cross-day association similarity and SCAD control

To quantify how stable association-specific population patterns were across days, we computed cross-day similarities for association-preferring neurons. For a given association and window, we took the trial-averaged ΔF/F time course within that window for each neuron and computed Pearson correlations between response vectors from different days. Correlations were Fisher *z*-transformed, averaged, and then back-transformed to yield cross-day similarity scores. We computed the same measure for SCAD neurons (cells tracked across days), which isolates per-cell pattern stability.

Statistics

Unless otherwise noted, α = 0.05 (two-tailed). We used Kruskal-Wallis tests for across-session effects, Fisher’s exact tests (and Fisher-Freeman-Halton Monte Carlo tests where appropriate) for contingency tables, Wilcoxon signed-rank tests for paired comparisons, Mann-Whitney rank-sum tests for unpaired comparisons, and exact binomial tests for deviations from 50% pathway composition. For multiple related comparisons we controlled for false discovery rate using the Benjamini-Hochberg procedure. For decoder learning trends we used linear mixed-effects models with day as a fixed effect and animal as a random intercept. For GLM coefficients, we used two-way ANOVA (Pathway × Session) on neuron-level β values.

## Acknowledgments

We thank Dr. Edgar Díaz-Hernández, Dr. Kasia Bieszczad, Dr. Todd Mowery, Dr. Rafiq Huda, and Dr. Marc Fuccillo for their valuable advice on experimental design, analysis, and conceptual development of this project; Dr. Alex Yonk and Sinduja Baskar for support with the animal colony and technical assistance; and members of the Margolis lab for helpful discussions.

## Funding

This work was supported by grants from the National Institutes of Health, (R01-NS094450 to D.J.M.) and National Science Foundation (IOS-1845355 to D.J.M.). I.L-G. was supported by a Rutgers Busch Biomedical Research grant and a Gordon and Betty Moore Foundation grant.

## Supplementary Material

**Supplementary Figure 1.**
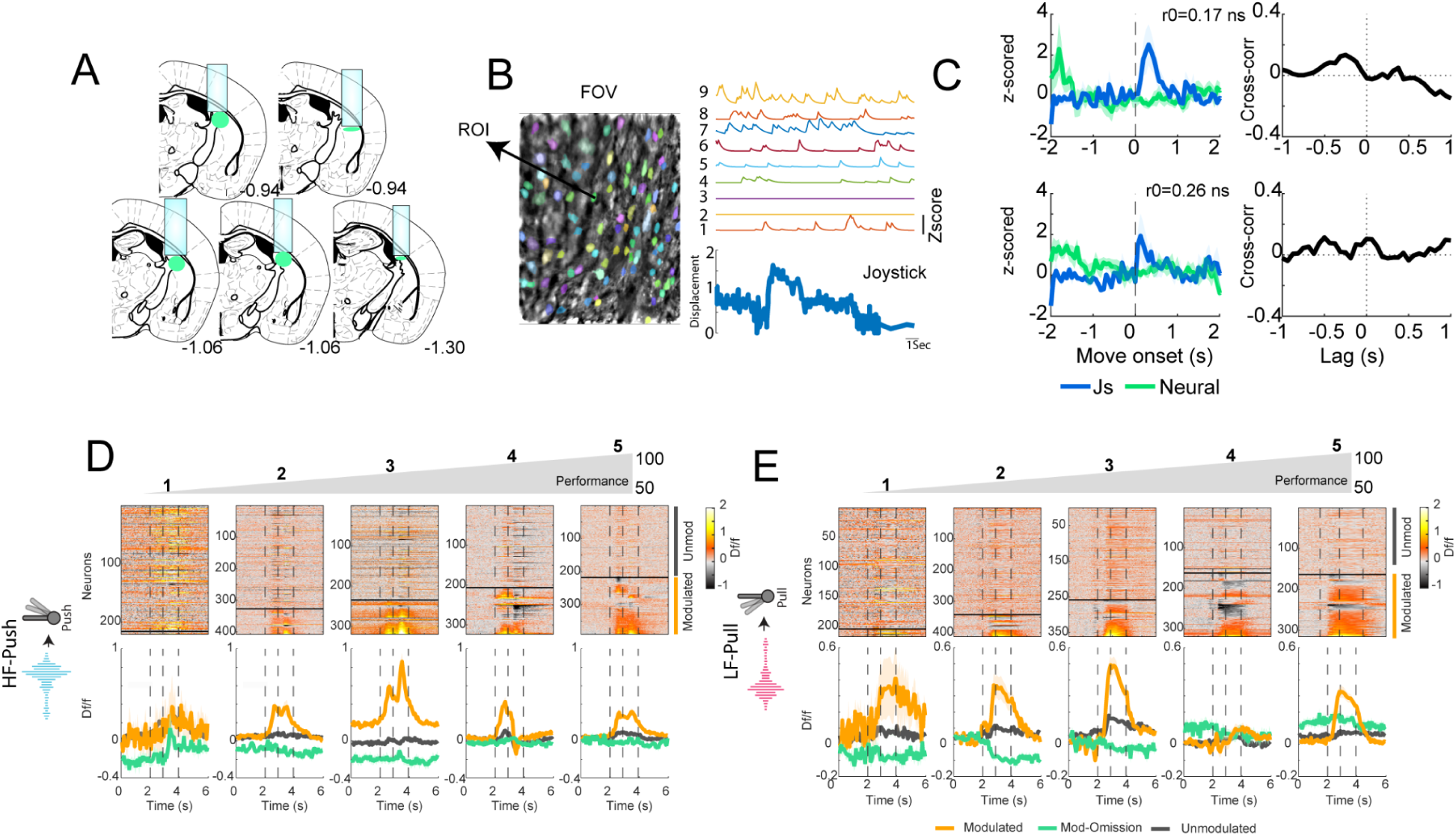
Recording sites, example imaging field, and controls for movement-related population activity. **A,** Recording sites across five mice (three D1-Cre × Ai14 and two A2a-Cre × Ai14) targeting TS (schematized reconstruction;Target AP −1.2 mm and ML 2.6 mm, left hemisphere). **B,** Example two-photon field of view and Suite2p segmentation output (ROIs) from a TS recording, with representative calcium traces shown alongside joystick displacement for the same session. **C,** Population activity aligned to joystick movement onset. For each session, joystick magnitude and mean population activity were resampled to a common timeline (−2 to +4 s around movement onset), z-scored, and compared using the zero-lag Pearson correlation (r₀) and the full cross-correlogram. Zero-lag coupling was small and not significant for push top (median r₀ = 0.053, IQR = 0.377; n = 5; Wilcoxon signed-rank vs 0, p = 0.594) and for pull down(median r₀ = 0.209, IQR = 0.074; n = 5; Wilcoxon signed-rank vs 0, p = 0.156). Cross-correlation showed modest off-zero peaks (push: median peak r = 0.299 at −1.0 s; pull: median peak r = 0.279 at +0.438 s), consistent with TS activity not being dominated by a unitary motor command. **D,** HF-Push across five days (low to high performance). Heatmaps: non-task modulated neurons (top) and task-modulated neurons (bottom) on rewarded trials. Mean traces (bottom): task-modulated rewarded, non–task-modulated rewarded, and task-modulated omission. **E,** Same as D for the LF-Pull association.

**Supplementary Figure 2.**
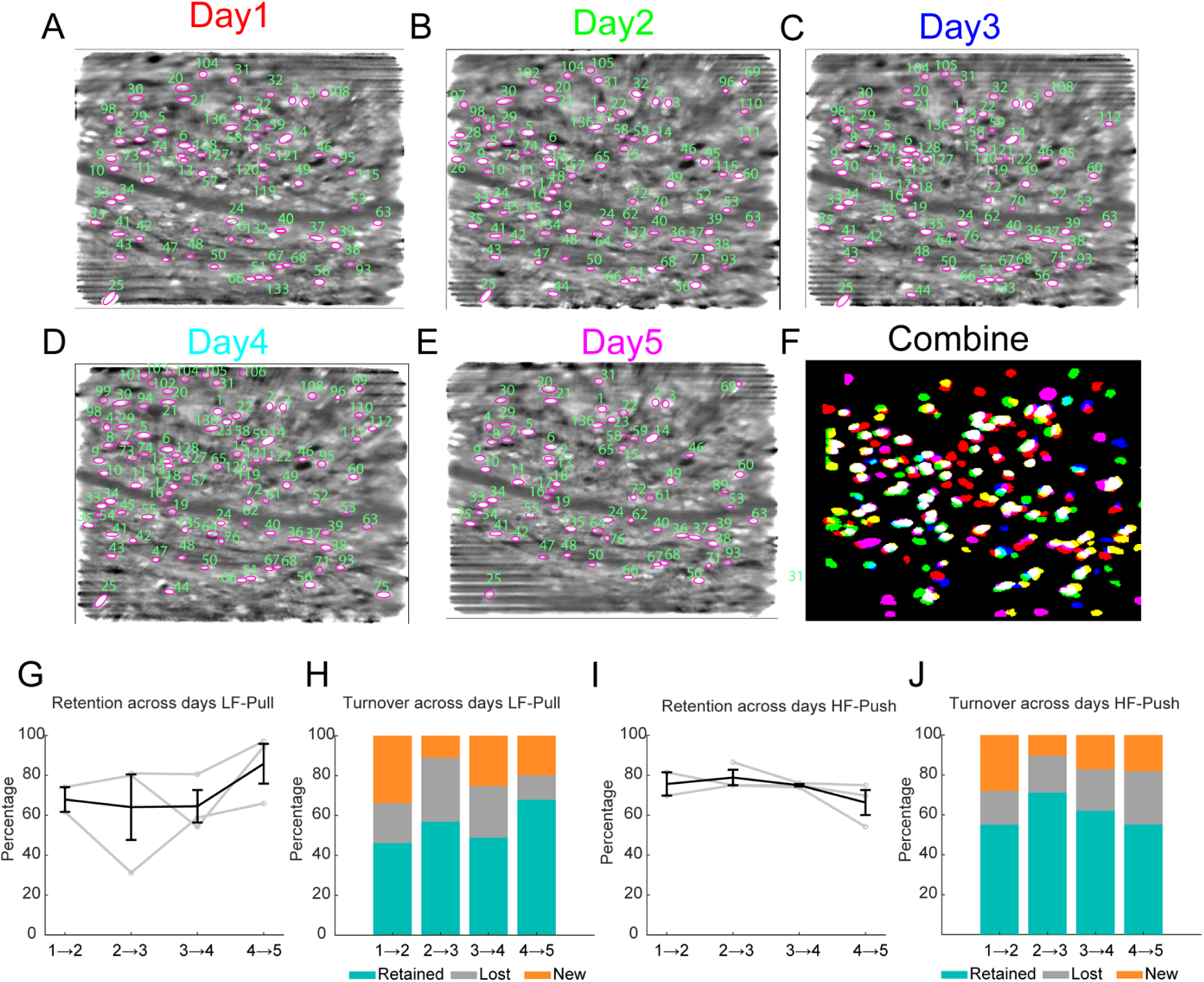
Cross-day ROI tracking (SCAD) and cell retention across performance days. **A-E,** Example mouse field of view across five recording days (Day 1-Day 5), showing Suite2p ROIs and cross-day matches. Each session was processed independently in Suite2p, and cross-day correspondences were established offline by overlaying session mean images and ROI masks and matching ROIs based on centroid proximity and contour/shape similarity. Matched ROIs were assigned a persistent neuron identifier (SCAD), whereas unmatched ROIs received new IDs. **F,** Combined ROI map across the five non-continuous recording days. ROIs are color-coded by day (Day 1 red, Day 2 green, Day 3 blue, Day 4 cyan, Day 5 magenta). White ROIs indicate overlap of matched ROIs across days (same cells identified in multiple sessions). **G,** LF-Pull. Percentage of retained cells across performance days. **H,** LF-Pull. Turnover summary across days, showing the fraction of cells retained, lost, and newly observed across performance days. **I,** HF-Push. Percentage of retained cells across performance days. **J,** HF-Push. Turnover summary across days, showing the fraction of cells retained, lost, and newly observed across performance days. Across the five representative performance days, retention was on the order of ∼40-60% in both associations.

**Supplementary Figure 3.**
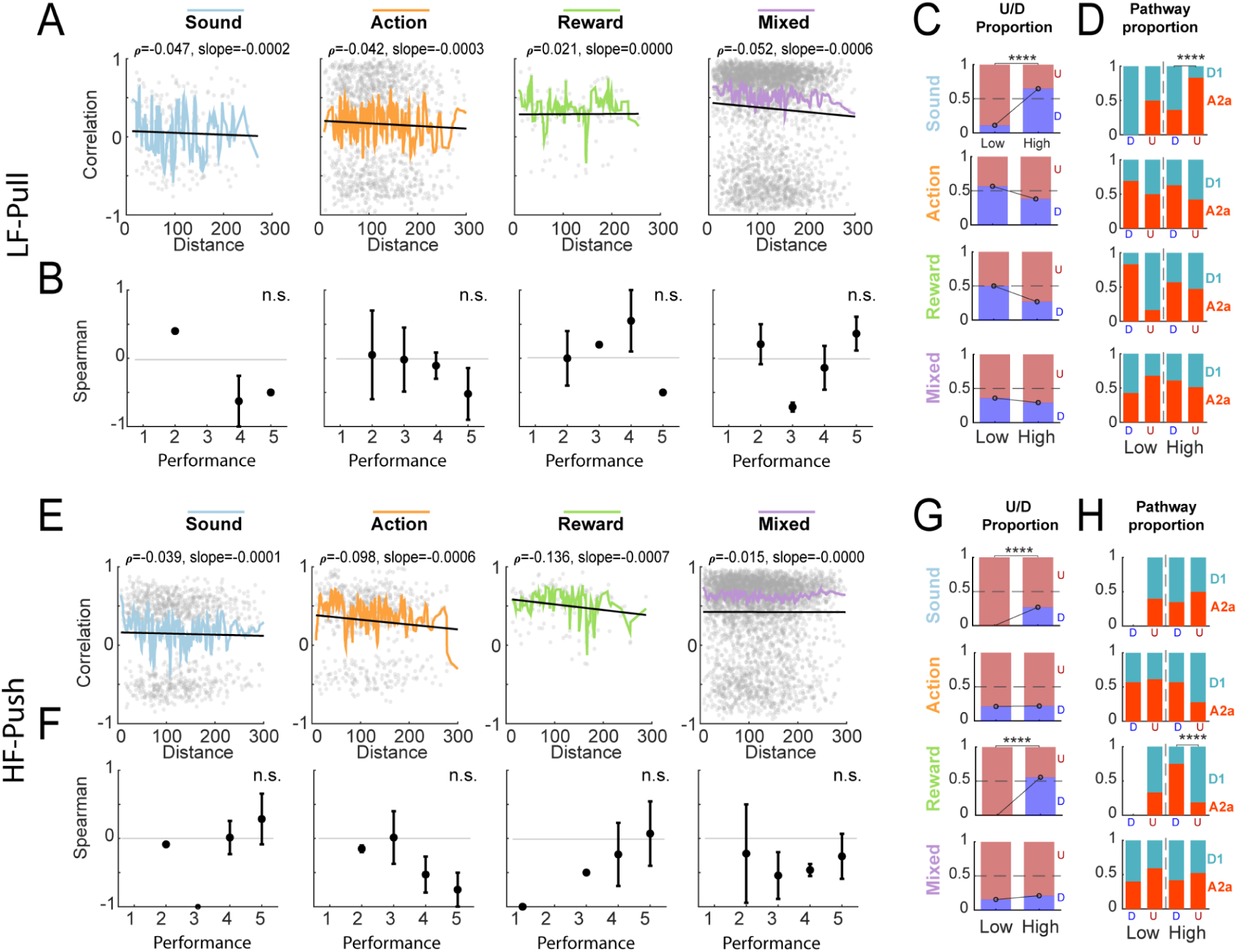
Up- and down-modulated category neurons across performance and lack of distance-similarity structure. **A,** LF-Pull. Pairwise functional similarity versus intersomatic distance within each category, showing no reliable distance-similarity relationship (Spearman; all p > 0.05). **B,** LF-Pull. Spearman ρ values for the distance-similarity relationship across performance days, showing no change across learning (Kruskal-Wallis across sessions; all p > 0.05). **C,** LF-Pull. Proportions of up- and down-modulated neurons on low- versus high-performance days within each category, showing an increase in down-modulated Sound neurons at high performance (Fisher’s exact test, p = 0.019). **D,** LF-Pull. Pathway composition (D1 vs A2a) for up- and down-modulated neurons within categories. A pathway difference is observed for Sound neurons at high performance, with a higher proportion of A2a cells in the down-modulated population (p = 0.0129). **E,** HF-Push. Pairwise functional similarity versus intersomatic distance within each category, showing no reliable distance-similarity relationship (Spearman; all p > 0.05). **F,** HF-Push. Spearman ρ values for the distance-similarity relationship across performance days, showing no change across learning (Kruskal-Wallis across sessions; all p > 0.05). **G,** HF-Push. Proportions of up- and down-modulated neurons on low- versus high-performance days within each category, showing increases in down-modulated Sound neurons (p = 0.0069) and down-modulated Reward neurons at high performance (Fisher’s exact tests). **H,** HF-Push. Pathway composition (D1 vs A2a) for up- and down-modulated neurons within categories. A pathway difference is observed for Reward neurons, with a higher proportion of D1 cells in the up-modulated population (p = 0.0020).

**Supplementary Figure 4.**
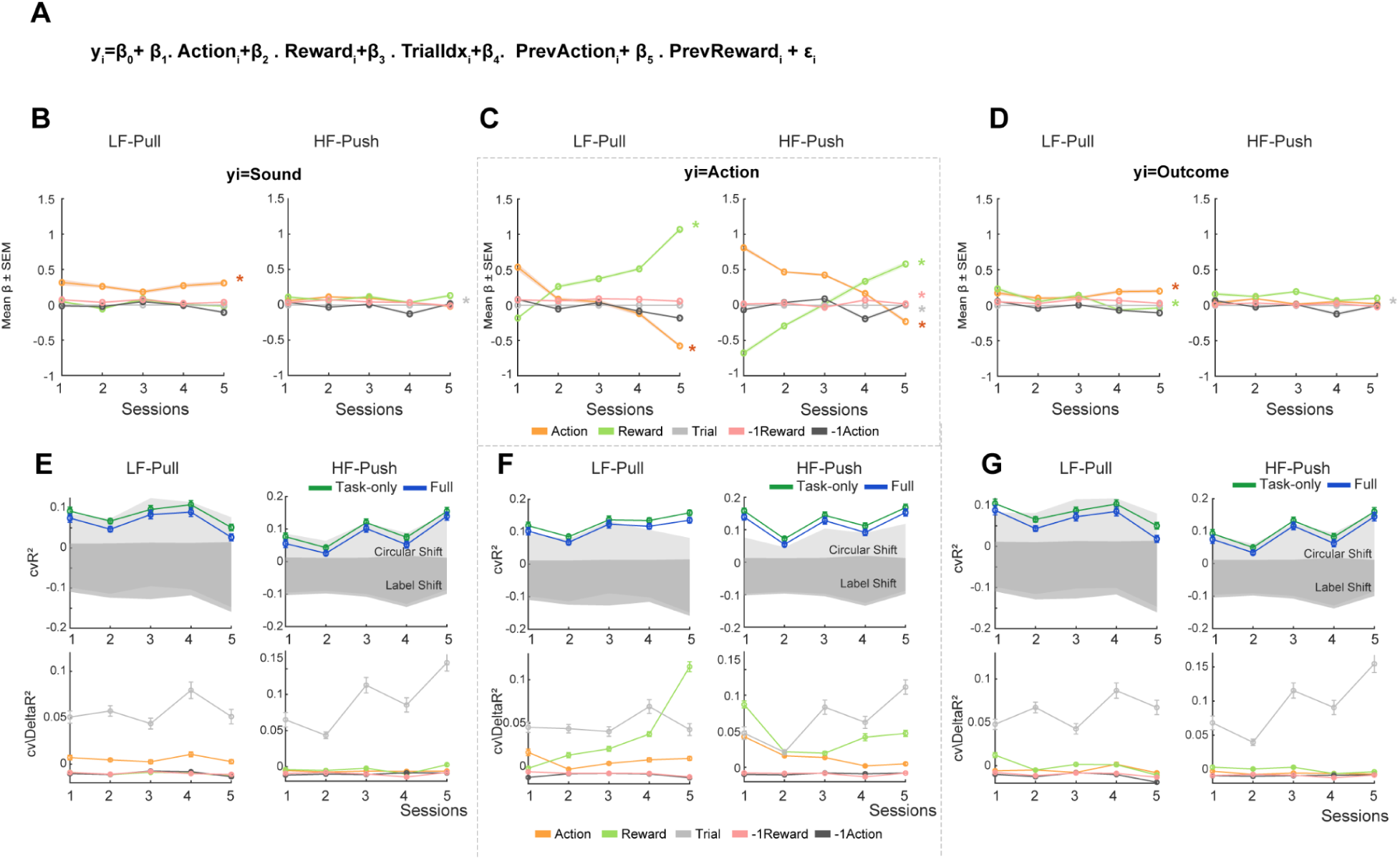
GLM prediction is strongest in the Action window and history terms contribute minimally. **A**, GLM structure. Trial-wise peak ΔF/F was modeled within Sound, Action, and Outcome windows using predictors for current-trial Action (movement vs omission), Reward (delivered vs not), and TrialIdx, with optional history terms (PrevAction and PrevReward). **B**, Sound window β coefficients across five representative sessions for LF-Pull (left) and HF-Push (right), shown for Action, Reward, TrialIdx, PrevAction, and PrevReward. **C**, Action window β coefficients across sessions for LF-Pull (left) and HF-Push (right), showing the learning-dependent increase in Reward β and decrease in Action β. **D**, Outcome window β coefficients across sessions for LF-Pull (left) and HF-Push (right), shown for the same predictors. **E**, Sound window model validation. Top, Task-only cvR² compared against two within-day null bands: a circular-shift null (preserves slow drifts but breaks trial timing) and a label-shuffle null (destroys trial structure). Bottom, unique contributions (cvΔR²) show minimal explained variance beyond TrialIdx. **F**, Action window model validation. Top, Task-only cvR² lies above both null bands for LF-Pull (left) and HF-Push (right), indicating reliable prediction of trial-by-trial peaks in the action epoch. Bottom, unique contributions (cvΔR²) show that Reward and Action account for most of the explained variance (with TrialIdx contributing more modestly). **G**, Outcome window model validation. Top, Task-only cvR² remains near the null bands for LF-Pull (left) and HF-Push (right). Bottom, unique contributions (cvΔR²) indicate minimal explained variance, dominated by TrialIdx.

**Supplementary Figure 5.**
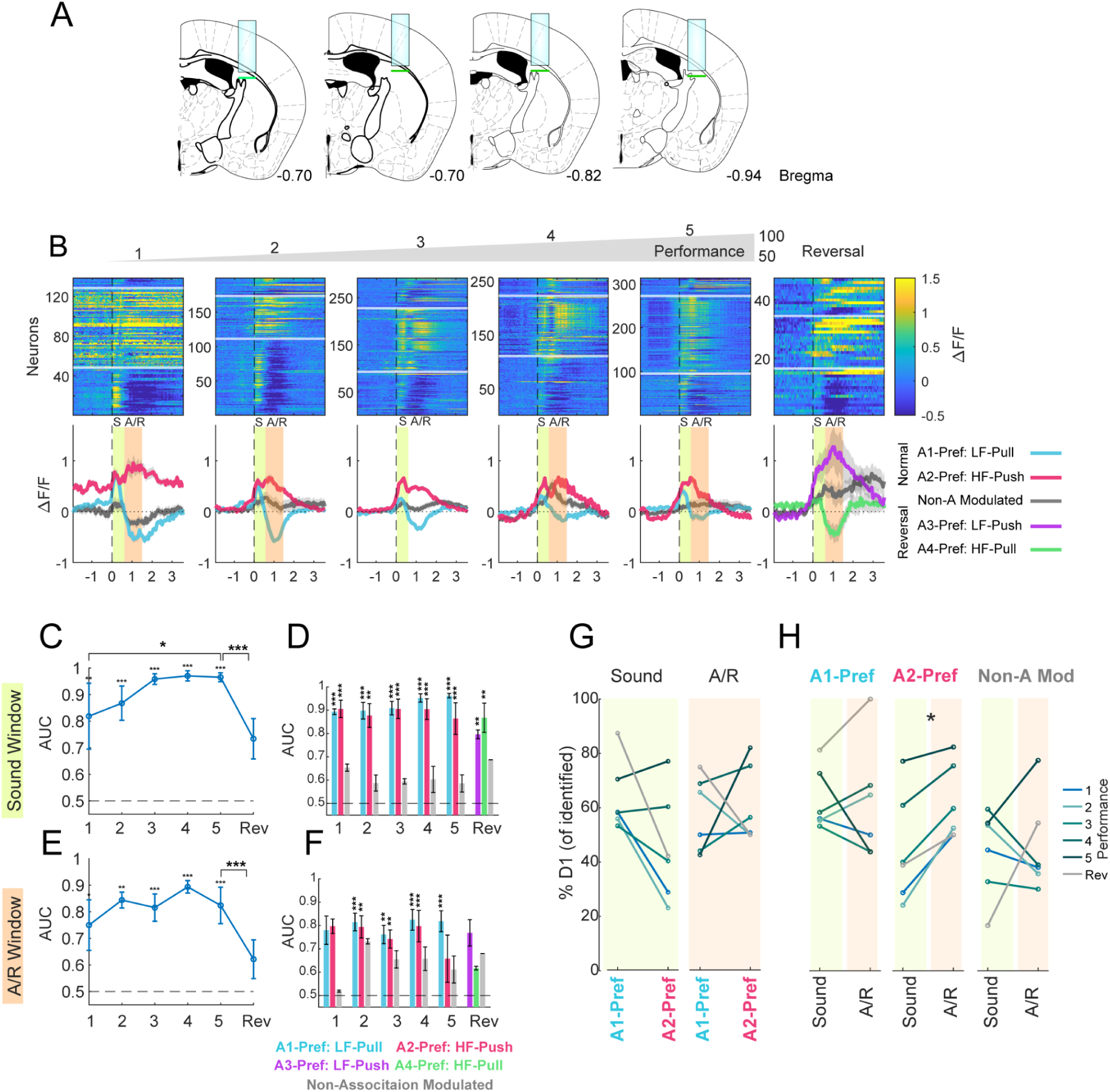
Association-preferring activity increases with learning and shows minimal pathway bias. **A**, Recording sites for the two-choice (2AFC) task. Sites from four mice (three D1-Cre × Ai14 and one A2a-Cre × Ai14) spanned ∼AP −0.70 to −0.94 mm from bregma in TS and were imaged through 1-mm GRIN lenses. **B,** Association selectivity across performance. TS SPNs were classified as A1-Pref., A2-Pref., or Non-A Mod. using an association-selectivity auROC computed on correct trials, where A1 = LF-Pull and A2 = HF-Push. In the reversal session, the sound-action associations were LF-Push and HF-Pull. **C,** Sound-window decoding. A decoder trained on sound-locked responses classified A1 versus A2 above shuffle on most days, improved with performance, and declined after reversal. **D,** Sound-window decoding across neuron groups. Decoding was above shuffle across performance days for association-preferring neurons and in the reversal session, but not for Non-A Mod. neurons. **E,** Action/reward-window decoding. Decoding remained above shuffle on most days but did not show a strong performance-dependent increase. **F,** Action/reward-window decoding across neuron groups. Decoding was above shuffle for association-preferring neurons on most days, but not for Non-A Mod. neurons, and declined in the reversal session. **G,** Pathway comparison for association preference. Pathway proportions were similar across association-preferring neurons in both the sound and action/reward windows. **H,** Window-by-pathway comparison. For A2-Pref. neurons, A2a cells were more prevalent in the sound window and D1 cells in the action/reward window (Wilcoxon signed-rank test, *p* < 0.05).

## Data Availability

https://github.com/margolislab/Linares-Garcia_2026

## Notes

### Competing Interest Statement

The authors have declared no competing interest.

